# Unveiling a unique macrophage population in exocrine glands sustained by ILC2-derived GM-CSF

**DOI:** 10.1101/2024.10.30.620897

**Authors:** Frederike Westermann, Selma Tuzlak, Victor Kreiner, David Bejarano, Mitchell Bijnen, Virginia Cecconi, Hannah van Hove, Haiting Wang, Gioana Litscher, Aline Ignacio, Rachel Lindemann, Laura Oberbichler, Donatella DeFeo, Zhaoyuan Liu, Anja Kipar, Kathy McCoy, Iain Nixon, Calum C Bain, Christoph Schneider, Sonia Tugues, Melanie Greter, Florent Ginhoux, Andreas Schlitzer, Elaine Emmerson, Burkhard Becher

## Abstract

Granulocyte-macrophage colony-stimulating factor (GM-CSF) has a non-redundant role in the emergence and maintenance of alveolar macrophages (AMs). However, its role in developmental and steady-state myelopoiesis outside the lung is largely unexplored.

Scanning through developing tissues using a Fate-map and reporter of GM-CSF mouse strain, we discovered that GM-CSF was produced by type 2 innate lymphoid cells (ILC2s) in the submandibular and sublingual salivary gland (SG) during postnatal development. GM-CSF producing ILC2s foster the development of a hitherto undescribed phagocyte subset, which we named adenophages. Detailed analysis focusing on phenotypic and transcriptional profiling revealed that adenophages display shared aspects of both, macrophages and dendritic cells (DCs). We found them to be homogenously distributed across the SG, but always in close proximity to GM-CSF producing ILC2s and myoepithelial cells. Importantly, adenophages were present throughout all analyzed exocrine glands such as lacrimal glands and mammary glands, and were also identified in human SG sections, indicating a conserved role in exocrine glands across species.

## Introduction

Granulocyte-macrophage colony-stimulating factor (GM-CSF, encoded by *Csf2*) was initially described as a colony-stimulating factor for both macrophages and neutrophils^1^. Whereas both macrophages and granulocytes have dedicated colony-stimulating factors – macrophage colony-stimulating factor (M-CSF, encoded by *Csf1*) and granulocyte colony-stimulating factor (G-CSF, encoded by *Csf3*), GM-CSF was found to stimulate the growth of dendritic cells (DCs) from bone marrow precursors *in vitro*^2^. *In vivo*, GM-CSF deficiency revealed a selective role of this cytokine for the emergence and maintenance of alveolar macrophages (AMs)^3–6^ and has been shown to support a subset of DCs located in barrier tissues^7–10^. Patients with autoantibodies against GM-CSF develop pulmonary alveolar proteinosis^11,12^, which revealed the importance of GM-CSF and AMs in processing surfactants^13,14^. Alveolar epithelial type II cells^4^ have been shown to be the non-redundant cellular source of GM-CSF needed for the survival of AMs. However, innate lymphoid cells (ILCs) can be an additional cellular source for GM-CSF under homeostatic conditions across multiple barrier tissues, where it is described to exhibiting explicit functions^10,15–18^. For example, GM-CSF derived from group 3 ILC (ILC3) in the intestine promotes antibacterial and immunomodulatory functions in intestinal myeloid cells^16,18^, while group 2 ILCs (ILC2s) were found to produce GM-CSF in the skin in order to maintain dermal cDC1s^10^. However, no other discernable defects in the macrophage, granulocyte and DC lineages were described in mice lacking GM-CSF. This is in line with the notion that GM-CSF is virtually absent under physiological conditions in the serum of mice and humans^19–21^. In contrast, mice and patients suffering from chronic inflammatory disease (e.g. rheumatoid arthritis, multiples sclerosis or sepsis) display a dramatic increase in GM-CSF stemming mainly from T cells and other lymphoid cells and closely correlating with disease activity^16,22–26^. This led to a shift in the research focus from the physiological role of GM-CSF towards its properties as a proinflammatory mediator across numerous inflammatory conditions, both pre-clinically and clinically^22,23,27–33^.

With the emergence of new research tools and the development of new transgenic mice, we revisited the role of GM-CSF in development and steady-state physiology outside of the lung. Combining GM-CSF fate mapping and gene targeting with single cell analysis to interrogate steady-state and development, we discovered a hitherto undescribed macrophage subset in exocrine glandular tissues during postnatal maturation, which is developmentally hardwired and depends specifically on ILC2-derived GM-CSF.

## Results

### ILC2s are a major source of GM-CSF in the postnatal salivary gland

To identify homeostatic GM-CSF production throughout mammalian postnatal development, we used the fate-map and reporter for GM-CSF mouse (FROGxAi14, Supplementary Figure 1A), which permanently labels GM-CSF producing cells and their progeny^23^. Comparing various organs, flow cytometry-based analysis revealed the salivary gland (SG, refers to submandibular and sublingual salivary glands) to harbor the highest frequency of GM-CSF producing cells within the hematopoietic compartment compared to other tissues analyzed including the lung (Figure 1A).

**Figure 1:**
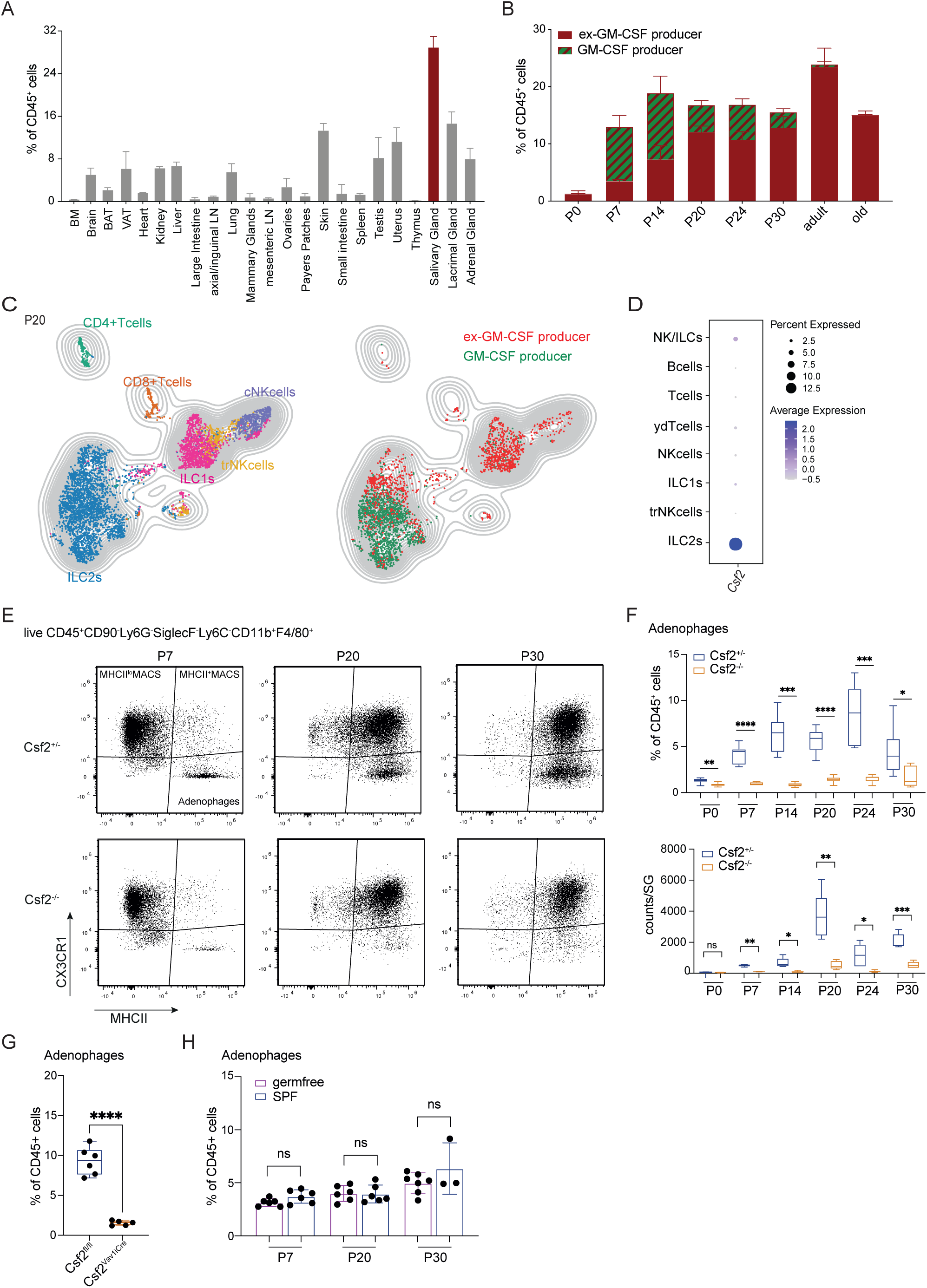
ILC2s produce GM-CSF in the postnatal salivary gland that harbors a GM-CSF dependent phagocyte population. A) Frequency of tdTomato^+^CD45^+^ cells in multiple organs of FROGxAi14 mice. Data are represented as mean±SD. B) Frequency of tdTomato^+^ (red=ex-GM-CSF producer) and tdTomato^+^eGFP^+^ (red/green = GM-CSF producer) CD45^+^ cells in the SG at indicated time-points. Data shows two independent experiments per timepoint and the data was pre-gated on singlets and live cells. Data are represented as mean±SD. C) Unsupervised clustering of HDCyto data focusing on the lymphoid compartment in the SG of FROGxAi14 mice at P20. Pre-gated on live CD45^+^CD11b^−^CD11c^−^CD19^−^ cells. Left UMAP shows identified cell subsets, right UMAP shows overlayed tdTomato^+^ (ex-GM-CSF producer) and tdTomato^+^eGFP^+^ (GM-CSF producer) cells across the identified cell subsets. For visualization all 6100 cells were used. D) *Csf2* expression in scRNAseq data of lymphocytes in the SG at P20. Colors indicate the average expression, circle sizes represent percentage of cells expressing the gene. E) Representative FACS plots for the expression of CX3CR1 and MHCII gated on macrophages (singlets live CD45^+^CD90^−^Ly6G^−^SiglecF^−^Ly6C^−^ CD11b^+^F4/80^+^) in the SG of *Csf2*^+/−^ and *Csf2*^−/−^ mice at P7, P20 and P30. F) Frequencies (top) and total counts (bottom) of adenophages throughout postnatal development in the SG of *Csf2*^+/−^ (blue) and *Csf2*^−/−^ (orange) mice. Pre-gated on live CD45^+^Lin^−^Ly6G^−^SiglecF^−^Ly6C^−^ CD11b^+^F4/80^+^ cells. Statistical test: 2-way ANOVA. n = 6-12 mice per group and timepoint from 2-3 independent experiments. G) Frequencies of adenophages in the SG of *Csf2*^fl/fl^ and *Csf2*^Vav1iCre^ mice at P20. Pre-gated on live CD45^+^Lin^−^Ly6G^−^SiglecF^−^Ly6C^−^CD11b^+^F4/80^+^ cells. Statistical test: t-test. Data is from 2 independent experiments. H) Frequency of adenophages in the SG of germfree (purple) and SPF (blue) mice at indicated time-points. Statistical test: 2-way ANOVA. Data is from 2 independent experiments per timepoint. Data are represented as mean±SD. P = postnatal day; ILC = Innate-like lymphocyte; cNK = conventional NK cells; trNK = tissue resident NK cells; SPF = specific pathogen free; ns = not significant

Analysis of adult FROGxAi14 mice revealed that about 30% of total CD45^+^ cells in the SG were tdTomato^+^ indicating that they have produced GM-CSF previously, hereafter termed ex-GM-CSF producers (Figure 1A&B). To determine the timepoints during development when GM-CSF was initially produced (which is indicated by GFP expression), we analyzed the SG at different stages of postnatal development. We noted a high frequency of GM-CSF production in the SG particularly between postnatal day 7 (P7) and P30 (Figure 1B, Supplementary Figure 1B). We applied high dimensional spectral flow cytometry (HDcyto) analysis on cells isolated from the SG of FROGxAi14 mice at P20 and observed that most innate-like lymphocytes such as conventional natural killer (cNK) cells, tissue resident natural killer (trNK) cells, ILC1s and ILC2s had produced GM-CSF at some point in the past. Strikingly, the active source of GM-CSF at P20 were predominantly ILC2s (Figure 1C, Supplementary Figure 1C&D). This finding was confirmed by single cell RNA sequencing (scRNAseq) of all CD45+ cells isolated from the SG at P20, showing *Csf2* expression almost exclusively within the ILC2 population (Figure 1D). To address whether non-hematopoietic cells also expressed *Csf2* at this timepoint, we performed single nuclei RNA sequencing (snRNAseq) on nuclei isolated from the total SG at P20. Opposed to *Csf1,* we did not observe significant expression of *Csf2* in the non-hematopoietic compartment in the SG (Supplementary Figure 1E) confirming ILC2s as the main source of GM-CSF in the SG at P20.

### Identification of a GM-CSF dependent macrophage subset in the developing salivary gland

GM-CSFR expression is abundant across myeloid cells and macrophages are the most prominent hematopoietic cells in the SG. Hence, the high GM-CSF production in the SG during postnatal development prompted us to investigate the responding cell subsets. Tissue-resident macrophages in the SG are characterized by their expression of the surface molecules CD11c and MHCII^34,35^. Using HDcyto we analyzed macrophages in the SG of GM-CSF-deficient (*Csf2*^−/−^) and control (*Csf2*^+/−^) mice during postnatal development (P0, P7, P14, P20, P24, P30). We found the two previously described subsets of macrophages, namely CX3CR1^+^MHCII^lo^ macrophages (MHCII^lo^ MACS), and CX3CR1^+^MHCII^+^ macrophages (MHCII^+^ MACS). As shown before^34^, while MHCII^lo^ MACS decreased over time, MHCII^+^ MACS increased and made up 30-40% of CD45^+^ cells by P30 (Supplementary Figure 1F). In addition to these canonical SG macrophages, we identified a CD11b^+^F4/80^+^Ly6C^−^CX3CR1^−^MHCII^+^ subset, which has not been described before (Figure 1E). Importantly, in *Csf2^−/−^* mice, we found this CX3CR1^−^MHCII^+^ subset to be absent, demonstrating its dependence on GM-CSF (Figure 1E&F). Hereafter we termed this cell type adenophages [Greek: gland-eater]. Adenophages were consistently absent in *Csf2*^−/−^ SGs (Figure 1F), while the two subsets of canonical macrophages (MHCII^lo^ MACS and MHCII^+^ MACS) were not altered, and neither were DCs (Supplementary Figure 1F&G). Throughout development, adenophages increased gradually, reaching their maximum numbers at P20 (Figure 1F).

As we could detect GM-CSF production only in CD45^+^ cells, we aimed to determine whether immune cells are a non-redundant source of GM-CSF for the emergence of adenophages. Therefore, we used *Vav1^iCre^* ^36^ mediated depletion of *Csf2* (*Csf2^Vav1iCre^*), which led to the loss of GM-CSF throughout the hematopoietic compartment. These mice phenocopied the loss of adenophages observed in *Csf2^−/^*^−^ mice demonstrating their dependence on leukocyte-derived GM-CSF (Figure 1G). Importantly, and as expected, AMs were unaffected in these mice (Supplementary Figure 1H), as they are exclusively dependent on alveolar epithelial type II cells producing GM-CSF^4^.

The timepoint of adenophage expansion (P7-P20) correlates with changes in the microbial colonization of the oral cavity^37,38^. Postnatal microbial colonization induces a host-response culminating in the tailored development of the local mucosal immune system^37,39^ and can influence the differentiation of phagocyte populations within mucosal tissues^40,41^. To determine whether the emergence of adenophages is initiated through microbial changes in the oral cavity, we analyzed the dynamics of adenophages during SG development in germfree mice. We could show that adenophages developed normally in germfree mice, demonstrating that their arrival is independent of the microbiome and thus is developmentally hardwired (Figure 1H, Supplementary Figure 1I).

Taken together, we discovered that the developing SG harbors lymphocytes (mainly ILC2s) which actively produce GM-CSF in the absence of inflammatory stimuli. Thereby, they provide a niche for a hitherto undescribed phagocyte population, which we termed adenophages.

### Adenophages have a unique transcriptional profile and are seeded by fetal monocytes

To better understand the nature of adenophages, we performed HDcyto focusing on myeloid cell populations in the SG of 20 days old mice (Figure 2A). We observed that adenophages expressed *bona-fide* macrophage markers like F4/80 and CD64 but also shared characteristic features with conventional DC2s (cDC2s) (e.g. CD209a) (Figure 2B). Furthermore, we identified the costimulatory receptor CD226 (*DNAM1*) as discriminating antigen on adenophages within the CD11b^+^ SG phagocytes (Figure 2B).

**Figure 2:**
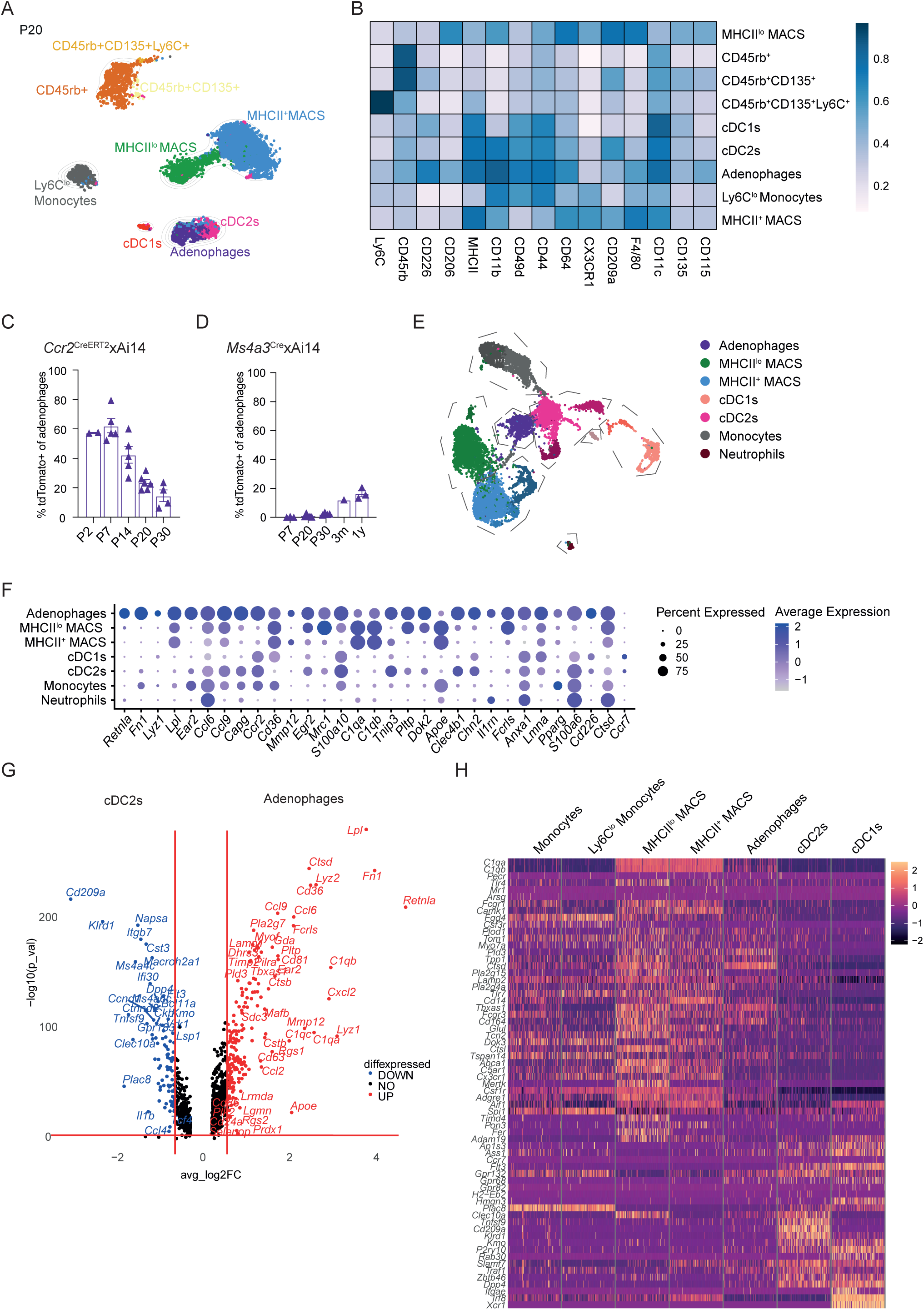
Adenophages show a distinct signature from other myeloid cells and arise from fetal monocytes. A) Unsupervised clustering of HDCyto data focusing on the myeloid compartment in the SG of Csf2^+/−^ mice at P20. Pre-gated on live CD45^+^CD90^−^SiglecF^−^Ly6G^−^ cells. For visualization in this UMAP 9461 cells were used. B) Heatmap of HDCyto data showing the marker expression within the myeloid cell populations defined in (A). C) Frequency of tdTomato^+^ adenophages in *Ccr2*^CreERT2^xAi14 mice labelled with tamoxifen at P0/P1. Bar plots show frequency at indicated postnatal timepoints. Data represents 1-2 independent experiments per timepoint. D) Frequency of tdTomato^+^ adenophages in *Ms4a3*^Cre^xAi14 mice. Bar plots show frequency at indicated postnatal timepoints. Data represents 1-2 independent experiments per timepoint. For C&D) Gating strategy for adenophages: singlets live CD45^+^CD90^−^Ly6G^−^SiglecF^−^Ly6C^−^CD11b^+^F4/80^+^MHCII^+^CX3CR1^−^ CD226^+^. Data are represented as mean±SEM. E) Unsupervised clustering of scRNAseq data of total myeloid cells in the SG of *Csf2*^+/−^ mice at P20. F) Gene signatures of adenophages, MHCII^lo^ MACS, MHCII^+^ MACS, cDC1s, cDC2s, monocytes and neutrophils in the SG at P20. Colors indicate the average expression of each gene, circle sizes represent percentage of cells expressing the indicated gene. G) Volcano plot showing DEGs between cDC2s and adenophages. Horizontal red line indicates P-value = 0.05; vertical red lines indicate log2FC = 0.4 or −0.4. H) Heatmap showing expression of macrophage and DC signature genes in the scRNAseq data of P20 SG (according to Immgen consortium 2012) of monocytes, MHCII^lo^ MACS, MHCII^+^MACS, adenophages, cDC2s and cDC1s in SG at P20.

Given that we could not clearly assign adenophages to a canonical macrophage or DC lineage, we aimed to better understand their origin using fate-mapping models for macrophage and DC development. First, we exploited the *Ccr2*^CreERT2^xAi14 strain^22^, which allows us to target fetal monocytes seeding the tissue in a perinatal wave when tamoxifen is applied at P0/P1 (Supplementary Figure 2A). Using this system, we observed that adenophages were of fetal monocytic origin (Figure 2C, Supplementary Figure 2B). Interestingly, the relative frequency of labelled cells decreased postnatally whereas their actual numbers remained unchanged (Supplementary Figure 2C). This suggests that the initially seeded fetal monocyte derived adenophages were not expanding but are accompanied by adenophages of postnatal origin. Most macrophages that arise postnatally are derived from granulocyte-monocyte-progenitors (GMPs)^42^. To address whether this is the case for adenophages, we analyzed the *Ms4a3*^Cre^xAi14 mouse model, which labels GMPs and their progeny. Surprisingly, adenophages were not derived from GMPs (Figure 2D), while GMP-dependent monocytes, neutrophils and eosinophils were effectively labelled in the *Ms4a3*^Cre^xAi14 (Supplementary Figure 2D). This suggests that adenophages are not canonical macrophages and are seeded by an alternative source postnatally as has previously been proposed^43^.

To better understand the molecular signature of adenophages, we performed scRNAseq on total CD45^+^ cells isolated from SG of *Csf2^+/−^* and *Csf2^−/−^* mice at P20, which resulted in 23 individual immune cell clusters (Supplementary Figure 2E). Focusing on the myeloid cell clusters we identified subsets corresponding to neutrophils (*S100a9, S100a8, Clec4d, Csf3r*), monocytes (*Plac8, Ly6c*), cDC1s (*Clec9a, Xcr1, Cd24a*), cDC2s (*Cd209a, Flt3, Irf4*), MHCII^lo^ MACS (*Mrc1, Lyve1, Folr2*), MHCII^+^ MACS (*Cx3cr, Hexb, Cd14*) and adenophages (*Retnla, Fn1, Cd226*) (Figure 2E&F, Supplementary Figure 2G). The canonical macrophages in the SG (MHCII^lo^ and MHCII^+^ MACS) correspond to conserved subsets of interstitial macrophages found across tissues^44^. MHCII^lo^ MACS expressed genes which are characteristic for macrophages that are juxtaposed to blood vessels^44^, while MHCII^+^ MACS resemble interstitial and duct-associated macrophages^44,45^. In contrast, the transcriptional signature of adenophages was clearly demarcated and unique, characterized by the expression of *Retnla (RELMα), Fn1, Mrc1, Egr2, Cd36* and *Cd226* (Figure 2F, Supplementary Figure 2F). The genes for the GM-CSF receptor (*Csf2ra*, *Csf2rb*) were abundantly expressed on all myeloid subsets similar to the M-CSF receptor (*Csf1r*) (Supplementary Figure 2G). However, the signature of adenophages profoundly featured genes associated with a GM-CSF imprinted signature, such as *Pparg, Cebpb* and *Irf4* (Supplementary Figure 2H)^22,27^. Interestingly, adenophages clustered close to cDC2s, in both, our HD-Cyto data as well as scRNAseq data (Figure 2A&E). To delineate as to whether adenophages and cDC2s share common transcriptional signatures, we analyzed the differentially expressed genes (DEGs) between adenophages and cDC2s in the SG at P20 (Figure 2G). cDC2s associated genes such as *Cd209a, Klrd1, Flt3, Ms4a6c* and *Plac8* were significantly lower expressed in adenophages. In contrast, adenophages were characterized by canonical macrophage genes such as *Rentla, Fn1, Cd36, Ccl6, Ccl9, Ear2* and *Mmp12* (Figure 2F&G, Supplementary Figure 2F). Comparing the adenophage signature to published macrophage and DC profiles from the Immgen consortium^46,47^, we found adenophages to predominantly express genes characteristic of macrophages (*Fn1, Mrc1, Fcgr1, Ctsd, Cd14, Csf1r, Adgre1, Lyz2*) (Figure 2H). Furthermore, we fed the signature of adenophages into the tabula muris data base and found it to predict a macrophage origin (Supplementary Figure 2I).

Taken together, adenophages are seeded by fetal monocytes and display a unique expression profile, which cannot be attributed to any hitherto described cell subset.

### Adenophages share morphological and functional characteristics with macrophages and DCs

Given the intermediate transcriptional features of adenophages, we isolated adenophages, MHCII^+^ MACS and DCs from the SG at P20 and imaged the cells by scanning electron microscopy (SEM). While the isolated macrophages and DCs were clearly identifiable as such, adenophages displayed a hybrid morphology, with cell membrane protrusions that were less pronounced than on cDCs (Figure 3A, Supplementary Figure 3A). However, when performing Giemsa stain on the same isolated subsets, we found adenophages to display morphological features common with macrophages, such as visible vacuoles (Supplementary Figure 3B).

**Figure 3:**
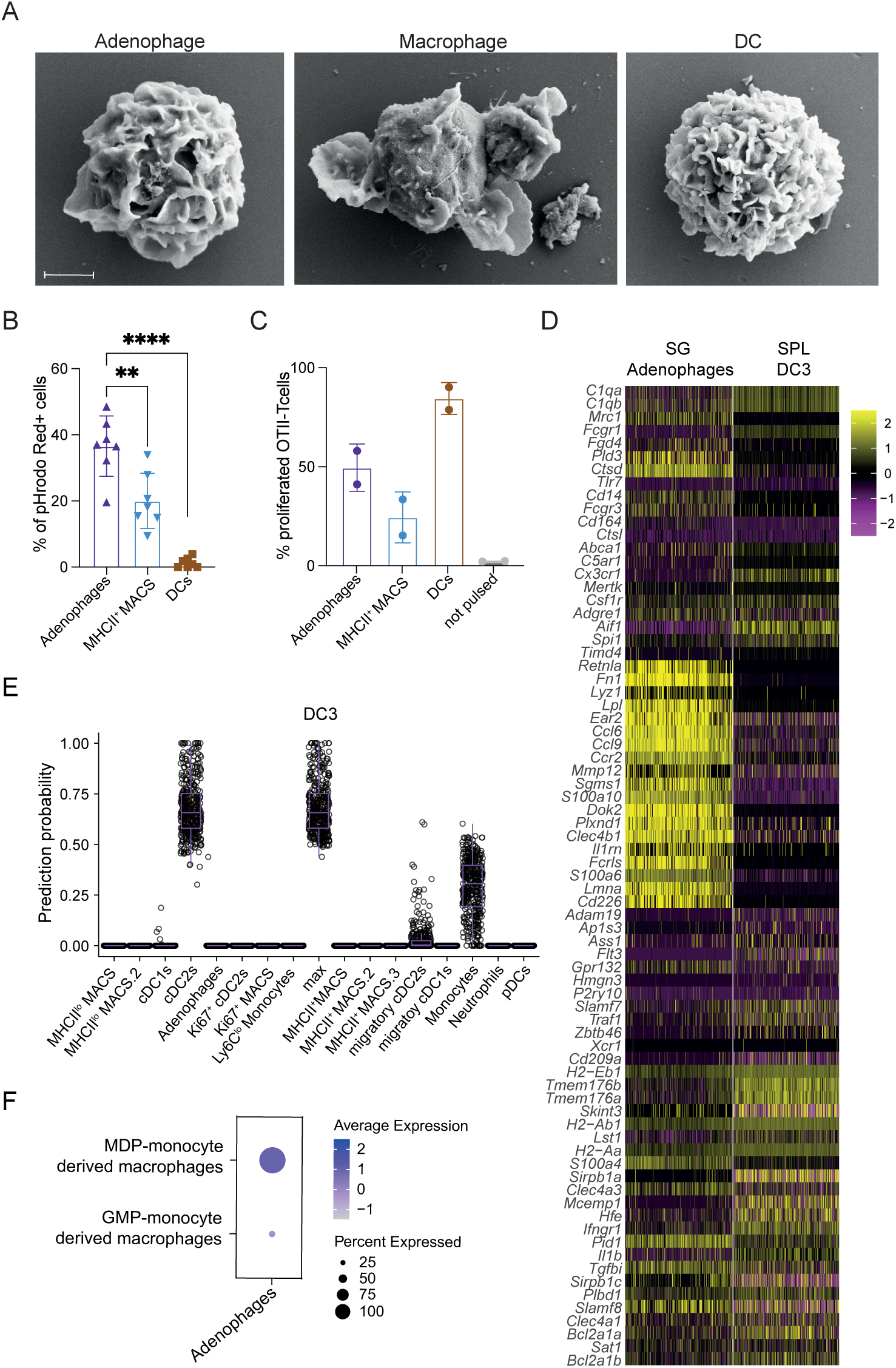
Adenophages share functional and morphological features with DCs and macrophages. A) Representative SEM images of adenophages, MHCII^+^MACS and DCs FACS-sorted from SG of Csf2^+/−^ mice at P20. Scale: 2µm. B) Frequency of pHrodo Red^+^ OT-II transgenic CD4^+^ T cells after 72 h co-culture with OVA_323–339_ pulsed adenophages (purple), MHCII^+^ MACS (blue), DCs (brown) or un-pulsed DCs (grey). The data represents two independent experiments. B&C) Data are represented as mean±SD. D) Comparison of gene expression in adenophages isolated from the SG and splenic DC3s. E) Probability prediction of DC3s falling into one of the myeloid clusters found in the SG. D-E) All analysis was performed on the integrated dataset of SG scRNAseq and splenic DC scRNAseq data (Liu et al., 2023). F) Gene signatures of MDP-monocyte derived macrophages or an GMP-monocyte derived macrophages found in adenophages. The signatures are from Trzebanski et al., 2024. Colors indicate the average expression of the gene signature, circle sizes represent percentage of cells within a cluster expressing the signature.

Macrophages and DCs are not only categorized by their ontogeny, but also by their functional properties. Hence, we studied adenophage function using *ex vivo* assays for phagocytosis and antigen presentation. Comparing the phagocytic capacity between adenophages and other SG resident myeloid cells, we found adenophages to be significantly more potent in phagocytosing *S.aureus* particles (Figure 3B, Supplementary Figure 3C), bacteria that are frequently associated with bacterial infections of the SG^48^. This indicates that adenophages are highly phagocytic and responsive to bacterial stimuli. Regarding their APC capacity, DCs are superior to adenophages in activating T cells (Figure 3C, Supplementary Figure 3D). However, adenophages outperformed other SG macrophage populations in their capacity to induce T cell proliferation, similar to what was described for dendritic cells type 3 (DC3s)^49^.

DC3s are a recently described subset originating from Ly6C^+^ MDPs and are presenting a monocyte/DC hybrid phenotype^49^. To determine as to whether adenophages describe SG DC3s, we integrated our scRNAseq dataset with the published DC3 scRNAseq data set^49^. The transcriptional signature of adenophages and DC3s is clearly demarcated, with adenophages prominently expressing genes such as *Retnla, Fn1, Ccr2, Mmp12, Ccl6, Ccl9* and *Cd226*, which we did not observe in DC3s (Figure 3D). Furthermore, probability prediction to evaluate whether DC3s or pro-DC3s describe any of the myeloid cells in our SG scRNAseq dataset did not show any overlap with adenophages (Figure 3E, Supplementary Figure 3E).

Although we excluded adenophages to be DC3s, the most likely postnatal source of adenophages were MDPs. Recently, Trzebanski *et al.* described the different molecular features of macrophages derived of MDP-monocytes compared to macrophages of GMP-monocyte origin^43^. Overlaying those gene signatures onto our scRNAseq data strongly suggested that MDP-monocytes contribute to the adenophage niche (Figure 3F).

Taken together, we show that adenophages have a hybrid morphology and are potently phagocytic and capable of T cell priming.

### Adenophages form a niche with GM-CSF producing ILC2s and myoepithelial cells

Upon recognizing the unique profile of adenophages, we were interested in their spatial distribution within the SG. Therefore, we performed Co-detection by Indexing (CODEX)-enabled high dimensional imaging^50^ on P20 SG of FROGxAi14 mice. Following a similar gating strategy as the one employed for flow cytometry, CODEX analysis allowed the identification of adenophages (and other hematopoietic and non-hematopoietic cell types) and revealed a homogenous distribution of adenophages across the SG (Supplementary Figure 4A). A cellular Neighborhood analysis centered on adenophages identified two tissue niches, primarily dominated by macrophages and with varying abundances of other myeloid and lymphoid cells, demonstrating that the adenophage niches in the tissue are rather homogenous (Supplementary Figure 4B). CODEX analysis also confirmed that the tdTomato signal as a surrogate marker for GM-CSF production co-localized with ST2^+^ ILC2s (Figure 4A). ILC2s were present in the cellular neighborhood of adenophages, and specifically tdTomato^+^ ILC2s were in close proximity to adenophages, when compared to GM-CSF-negative ILC2s (tdTomato^−^ ILC2s) (Figure 4B). This observation was confirmed through immunofluorescent staining for adenophages and GM-CSF producing cells in P20 SG of FROGxAi14 mice (Supplementary Figure 4C). Additionally, by interrogating our scRNA-seq data for cell-cell interactions using CellPhoneDB^51^ we observed a strong interaction between the GM-CSFR on adenophages and *Csf2* delivered by ILC2s (Supplementary Figure 4D).

**Figure 4:**
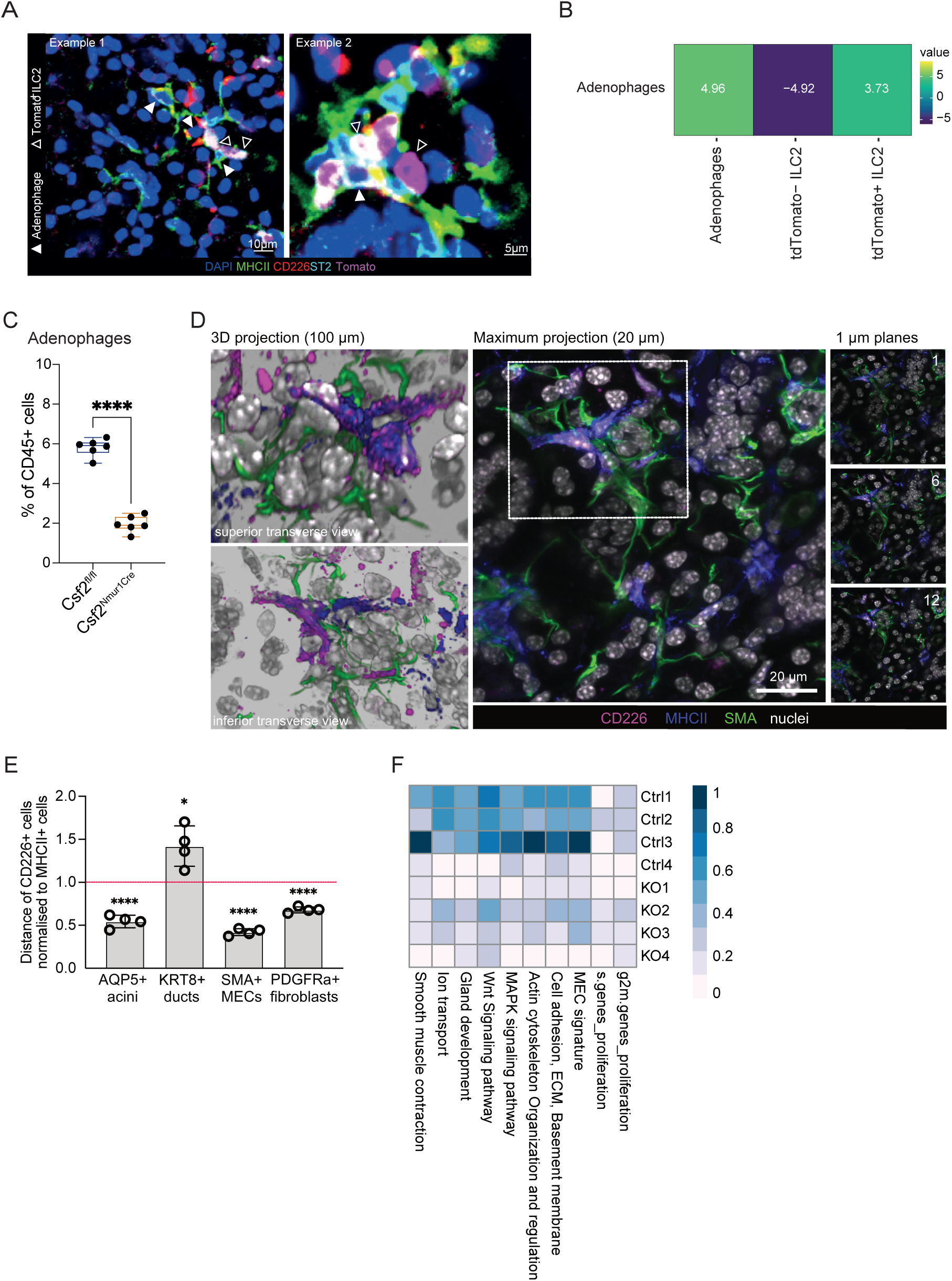
Adenophages form a niche with ILC2s and MECs. A) CODEX imaging showing indicate that cells have a positive co-localization C) Frequency of adenophages within CD45^+^ cells in *Csf2*^fl/fl^ and *Csf2*^Nmur1Cre^ (ILC2 specific deletion of GM-CSF) mice at P20. Statistical test: t-test. D) 3D projection of adenophages within the SG of mice at P20. Magenta = CD226, Blue= MHCII, Green = aSMA, white = DAPI. E) Quantification of the distance of adenophages from AQP5+ acini, KRT8+ ducts, SMA+ MECs and PDGFRa^+^ fibroblasts relative to other MHCII^+^ cells. Data are represented as mean±SEM. F) MEC signature indices in RNAseq samples from MECs of *Csf2*^+/−^ Control and *Csf2*^−/−^. Indices were calculated based on previously published gene signatures (Lu et al., 2012; Hauser et al., 2020; Mauduit et al., 2024; Thiemann et al, 2022). MEC = myoepithelial cell.

Above, we confirmed that adenophages rely on the delivery of GM-CSF through hematopoietic cells (Figure 1G). In order to definitively delineate whether ILC2s are the relevant source of GM-CSF for adenophage development, we generated *Nmur1*^Cre^xCsf2^fl/fl^ mice (*Csf2^Nmur1Cre^*)^52,53^. This specific deletion of GM-CSF only in ILC2s resulted in the loss of adenophages in the SG clearly demonstrating ILC2s as the non-redundant source of GM-CSF for the emergence of adenophages (Figure 4C).

Besides their interaction with other hematopoietic cells, we were also interested in whether adenophages interact with non-immune cells within the SG. We analyzed the scRNAseq dataset on murine SG from Hauser et al.^54^, which mainly focused on non-hematopoietic cells, allowing us to investigate cell-cell interactions between adenophages and stromal/epithelial cells. We applied CellChat, which revealed a particularly strong interaction of adenophages with mesenchymal, stromal and myoepithelial cells (MECs) (Supplementary Figure 4E). Specifically, the association with MECs was further strengthened by immunofluorescent histology showing superior spatial association of adenophages with MECs surrounding SG acini compared to other MHCII^+^ cells (Figure 4D&E, Supplementary Figure 4F). To explore the relationship of adenophages and MECs, we FACS-isolated MECs from SG of *Csf2 ^−/−^* and littermate control mice and performed RNAseq analysis. We confirmed that the isolated cells were indeed MECs as they expressed high levels of characteristic genes such as *Acta2* (α-SMA) and *Myh11* (Supplementary Figure 4G). Looking at DEGs we observed that MECs did not show drastic transcriptional differences in the absence of adenophages (Supplementary Figure 4H). However, when applying different gene signature scores, either for overall MEC identity or for described MEC functions (Supplementary Figure 4I, Supplementary Table 1^54–57)^, we observed a clear reduction of all identity/functionality scores in MECs isolated from SG devoid of adenophages, while their proliferation scores (Seurat Cell cycle scoring) were the same (Figure 4F, Supplementary Figure 4I). The quantifiable difference is not caused by significant DEGs but rather a consequence of slightly but consistently lower gene expressions of the signature genes described in Supplementary Table 1 (Supplementary Figure 4I).

Taken together, the niche of adenophages is characterized by the close proximity of GM-CSF producing ILC2s, which support their emergence, and MECs, which in turn are supported in their identity commitment by adenophages.

### Adenophages are conserved throughout multiple exocrine glands

The presence of adenophages in SG and their role in sustaining the MEC-identity prompted us to investigate whether they can also be found in other exocrine glands, which harbor MECs. Therefore, we analyzed the lacrimal gland (LG) and mammary gland (MG) of *Csf2*-proficient mice. Indeed, in both tissues we found a population expressing the same surface markers (MHCII^+^CX3CR1^−^CD226^+^) as SG-adenophages (Figure 5A&B, Supplementary Figure 5A). Importantly, adenophages were also absent from the LG and MG of *Csf2*^−/−^ mice to the same extent as observed for the SG (Figure 5C). Furthermore, we confirmed by IHC staining that adenophage associated with MECs in the LG (Supplementary Figure 5B). Analyzing published scRNAseq data on MG^58^, we were also able to identify an adenophage-like population in the naïve MG which is retained throughout lactation (Figure 5D, Supplementary Figure 5C). Importantly, in non-glandular tissues such as spleen, and skin we found only few cells expressing surface markers characteristic for adenophages (Supplementary Figure 5D). However, these cells were GM-CSF independent, arguing that they are not adenophages (Figure 5E).

**Figure 5:**
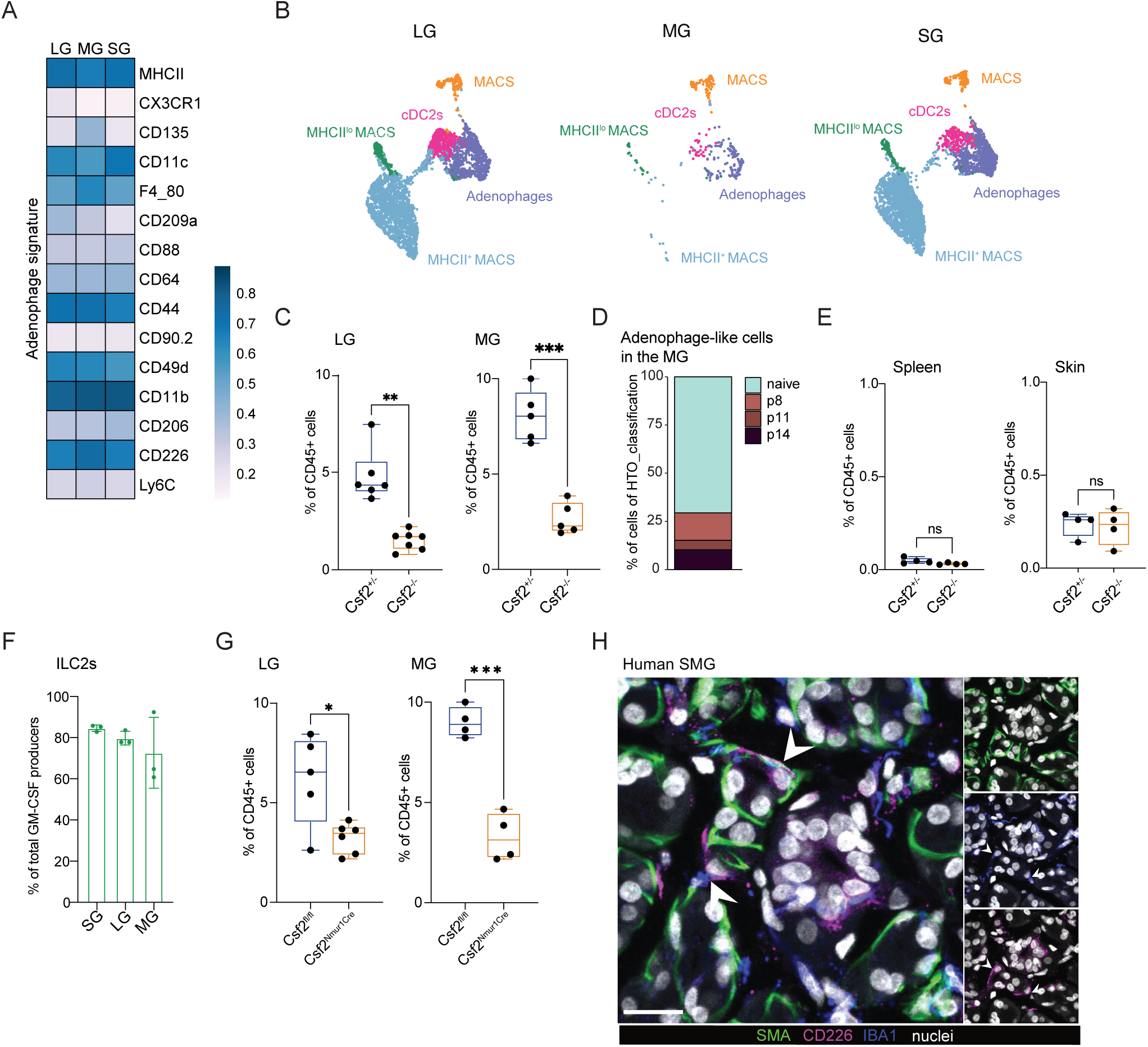
Adenophages are conserved across exocrine glands and species. A) Heatmap showing marker expression of adenophages in LG, MG and SG. B) Unsupervised clustering of HDCyto data focusing on the myeloid compartment in the LG, MG and SG of Csf2^+/−^ mice at P20. Pre-gated on live CD45^+^CD90^−^SiglecF^−^Ly6G^−^Ly6C^−^CD11b^+^ cells. For visualization in this UMAP all cells were used (LG: 4923, MG: 331, SG: 4445). C) Frequency of adenophages in the LG (left) and the MG (right) in Csf2^+/−^ and Csf2^−/−^ mice at P20. Statistical test: t-test. Data represents two independent experiments. D) Abundance of adenophage-like cells in a scRNAseq data set (Cansever, Petrova, et al. 2023) of naïve and lactating MG. E) Frequency of adenophage-like cells in the spleen (left) and the skin (right) in Csf2^+/−^ and Csf2^−/−^ mice at P20. Statistical test: t-test. F) Frequency of ILC2s within GM-CSF producing (tdTomato^+^GFP^+^) cells in the SG, LG and MG of FROGxAi14 mice at P20. Data are represented as mean±SD. Immunofluorescent histology staining of adenophages in human SMG. Scale = 20 µm. LG = lacrimal gland; MG = mammary gland; SMG = submandibular gland

Of note, also in MG and LG ILC2s were the main source of GM-CSF (Figure 5F, Supplementary Figure 5E). IHC staining confirmed that GM-CSF producing cells were closely associated with adenophages also in the LG (Supplementary Figure 5F). Depletion of ILC2 derived GM-CSF using *Csf2^Nmur1Cre^* mice resulted in the loss of adenophages in LG and MG, pointing towards a conserved ILC2-adenophage communication axis across exocrine glands (Figure 5G).

Lastly, we explored whether adenophages can be also found in humans. Therefore, we performed IHC staining for SMA (identifying MECs), IBA1 (a macrophage marker) and CD226 on human submandibular SG sections. Strikingly, we again found IBA1^+^CD226^+^ cells in close proximity to MECs also in human tissue, pointing towards a conserved cell type and cellular niche across species (Figure 5H).

Taken together, adenophages can be found throughout exocrine glands where they form a niche-triad with GM-CSF producing ILC2s and MECs.

## Discussion

Commonly, the GM-CSFR is expressed on DCs and their precursors as well as by monocytes and macrophages enabling all subsets to sense and respond to GM-CSF^4,5,7–9,32,59^. However, while GM-CSF has gained most attention for its role in inflammation over the past years^22,23,27^ only few myeloid cell subsets depend on its presence in the steady-state, the most prominent being AMs ^4,5,7–9,59^. Even though several cellular sources of GM-CSF have been described in the steady-state ^10,15,16,18^, within the lungs for instance the cellular source of GM-CSF required for the survival of AMs are exclusively alveolar epithelial type II cells^4^. The special relationship between alveolar epithelial type II cells and AMs forms the niche of the AM network dedicated to surfactant clearance.

Using sensitive fate map and reporter systems, we show that immune cells produce GM-CSF at some level in most analyzed tissues of the unchallenged murine organism, most prominently in the developing SG. The development of the SG starts early in the embryo and is not fully mature until puberty, constantly undergoing tissue rearrangement^54,60–62^. Dynamic changes in the immune compartment accompany the developmental restructuring of the SG, until it stabilizes in the adult animal. Throughout all developmental stages of the SG, macrophages are the predominant immune cell type. In the adult SG, tissue-resident macrophages have been shown to depend on *Csf1*^34,35^. Here, we discovered adenophages, a hitherto undescribed macrophage subset in the SG which depends on tonic GM-CSF production by ILC2s. ILCs have been described to produce GM-CSF in various barrier tissues, such as the intestine, lung, skin and SG^7,10,15–18^. We show that ILC2-derived GM-CSF is necessary and sufficient for the presence of adenophages specifically in postnatal exocrine glands, which is in stark contrast to the AM network in the lung. Consistently, adenophages are always found in close proximity to GM-CSF producing ILC2s.

Adenophages arise independently of the microbiome. Indeed, a recent study also described that the general immune compartment of the developing SG emerges independent from the microbial changes that coincide with postnatal development^37,39^. Although expressing markers identifying them as macrophages (e.g. F4/80 and CD64), adenophages show characteristics of both, macrophages and DCs. Consequently, we observed that adenophages were both, highly phagocytic as well as capable antigen-presenting cells via MHCII. The APC capacity of macrophages is generally inferior to those of DCs^63,64^. Of note, recently, Haimon et al. described productive APC interactions specifically between microglia and regulatory T cells^65^. Whether adenophage-APC properties are also dedicated to a specific T cell subset remains to be shown.

While the distinction of macrophages and DCs based on marker expression can be challenging due to overlapping expression profiles, their ontogeny can provide additional information to distinguish DCs from macrophages: tissue macrophages can originate during fetal myelopoiesis followed by a varying contribution of GMP-derived monocytes based on the respective tissue, while DCs arise from CDP progenitors^66,67^. Similar to the recently described DC3 subset^49^, the origin of adenophages also follows a non-classical ontogenetic path. While they are initially seeded by fetal monocytes in a perinatal wave, they are then accompanied by a second wave of adenophages postnatally. Through lineage tracing, they could not be assigned to be originating from GMPs. Recently, it has been shown that MDP-derived monocytes also contribute to peripheral tissue macrophage populations^43^, making them a likely postnatal source of adenophages. The strongly overlapping transcriptional signature to MDP-monocyte derived macrophages indicates a shared ontogeny with adenophages.

Regardless of their ontogeny, it is becoming increasingly clear that tissue-resident macrophages and DCs can adjust their characteristic transcriptional profile to adapt to the environmental niche e.g. Kupffer cells acquire a Lipid-associated macrophage (LAMs) signature in the context of liver injury and become LAM-like KCs ^68^. Hence, the characteristic phenotype and transcriptional signature of adenophages may well be shaped by GM-CSF imprinting and the microenvironment in the developing glands and by that overrule characteristics imprinted by ontogeny.

While *Csf1*-dependent SG macrophages have been shown to be involved in epithelial regeneration^35^, adenophages are closely associated with MECs surrounding the secretory cells of exocrine glands, the acini. MECs exhibit both epithelial and mesenchymal features and are essential in supporting the excretion of the glandular products^56,69^. Globally, adenophages did not drastically influence the transcriptional profile of MECs as we observed only select significantly DEGs. However, applying various gene signatures that have been described for MEC identity and function across exocrine glands^54–57^ it became apparent that the absence of adenophages led to reduced identity indices in MECs. This reduced MEC identity signature was not sufficient to cause any overt morphological differences in MECs in the absence of adenophages, however adenophages appear to be involved in maintaining a stable and fully differentiated and functional MEC population.

Importantly, although we first identified adenophages in the developing SG, we also found them in other exocrine glands such as LG and MG. This is in line with a recent report showing an adenophage-like population in a scRNAseq atlas of the LG (termed *Pf*^+^ M2 Mɸ)^70^. Similar to SG, in LG and MG ILC2-derived GM-CSF is essential for adenophages, strengthening the ILC2-adenophage axis. Importantly, although ILC2-derived GM-CSF in steady-state skin is described to support cDC homeostasis^10^, we did not find cells resembling adenophages in the skin, arguing for a glandular-specific niche of adenophages with ILC2s. The association of adenophages with MECs across exocrine glands points towards a conserved niche-triad of adenophages, ILC2s and MECs. Furthermore, we found that adenophages are present also in human SG, underlining the conserved nature of the adenophages niche in evolution.

Taken together, adenophages describe a novel phagocytic cell type that forms a network with GM-CSF producing ILC2s and MECs throughout developing exocrine glands and is conserved across species. It will be of great interest to further explore the role of adenophages in healthy glandular function but also under pathological conditions including degeneration, cancer and infection.

## Limitations of the study

In this study, we explored tissues from mice and human only. We believe it will be important to analyze adenophages across multiple species to understand the emergence of adenophages in vertebrate evolution. Furthermore, it is difficult to assess exocrine gland products during early murine development due to limited volume and ethical concerns regarding animal welfare. This study is essentially limited to juvenile animals in the steady-state focusing on adenophages in developing exocrine glands. In the future it will be important to study their role in adulthood and aging. Additionally, future studies to better understand the role of adenophages across various pathologies including degeneration, cancer and infection will be of great interest.

## Acknowledgments

We thank the UZH Center for Microscopy and Image Analysis for help with SEM; the Functional Genomics Center (University of Zurich) for technical support with the scRNAseq, snRNAseq and bulkRNAseq experiments, the UZH Laboratory Animal Service Center for animal mouse husbandry; We also would like to thank Lino Casty, Mirjam Lutz and Philipp Häne for technical support and Can Ulutekin for providing data analysis scripts.

The work was supported by funds from the European Research Council (ERC) under the European Union’s Horizon 2020 research and innovation program grant agreement no. 882424, and the Swiss National Science Foundation (170320, 188450, 219287, CRSII5_183478 and CRSII--222718).

## Author contributions

F.W., S.T., and B.B. conceived the study; F.W. and S.T. designed, performed, and analyzed most experiments, apart from specific contributions outlined here; V.K. performed computational data analysis of the sequencing experiments; D.B. and A.S. performed the CODEX experiments and analysis; E.E. performed the immunofluorescent immunology on mouse and human tissue; A.I. performed the adenophage analysis in germfree mice, M.B., V.C., H.vH., H.W., G.L., R.L. and L.O. supported mouse experiments. I.N. provided the human SMG samples. C.S., Z.L. and F.G. provided mice. B.B. supervised the study; F.W., S.T., and B.B. wrote the manuscript with contributions from F.G., E.E., and all other authors.

## Material and Methods

### Mouse strains

The mouse strains used in this study as following

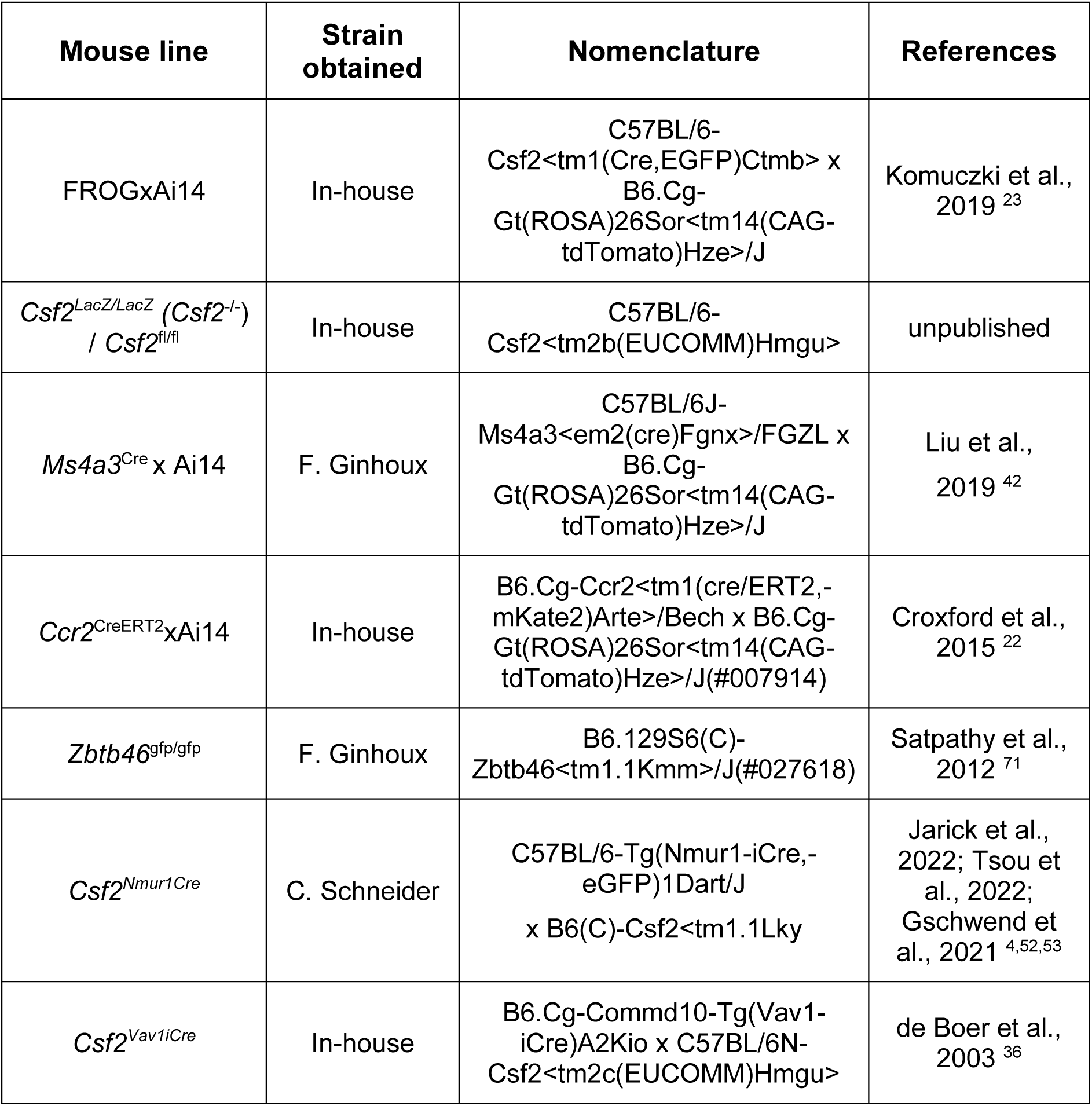

| Mouse line | Strain obtained | Nomenclature | References |
| --- | --- | --- | --- |
| FROGxAi14 | In-house | C57BL/6-Csf2<tm1(Cre,EGFP)Ctmb> x<br>B6.Cg-Gt(ROSA)26Sor<tm14(CAG-tdTomato)Hze>/J | Komuczki et al.,<br>2019 <sup>23</sup> |
| *Csf2*<sup>LacZ/LacZ</sup> (*Csf2*<sup>-/-</sup>)<br>/ *Csf2*<sup>fl/fl</sup> | In-house | C57BL/6-Csf2<tm2b(EUCOMM)Hmgu> | unpublished |
| *Ms4a3*<sup>Cre</sup> x Ai14 | F. Ginhoux | C57BL/6J-Ms4a3<em2(cre)Fgnx>/FGZL x<br>B6.Cg-Gt(ROSA)26Sor<tm14(CAG-tdTomato)Hze>/J | Liu et al.,<br>2019 <sup>42</sup> |
| *Ccr2*<sup>CreERT2</sup>xAi14 | In-house | B6.Cg-Ccr2<tm1(cre/ERT2,-<br>mKate2)Arte>/Bech x B6.Cg-Gt(ROSA)26Sor<tm14(CAG-tdTomato)Hze>/J(#007914) | Croxford et al.,<br>2015 <sup>22</sup> |
| *Zbtb46*<sup>gfp/gfp</sup> | F. Ginhoux | B6.129S6(C)-<br>Zbtb46<tm1.1Kmm>/J(#027618) | Satpathy et al.,<br>2012 <sup>71</sup> |
| *Csf2*<sup>Nmur1Cre</sup> | C. Schneider | C57BL/6-Tg(Nmur1-iCre,-<br>eGFP)1Dart/J<br>x B6(C)-Csf2<tm1.1Lky | Jarick et al.,<br>2022; Tsou et<br>al., 2022;<br>Gschwend et<br>al., 2021 <sup>4,52,53</sup> |
| *Csf2*<sup>Vav1iCre</sup> | In-house | B6.Cg-Commd10-Tg(Vav1-iCre)A2Kio x C57BL/6N-Csf2<tm2c(EUCOMM)Hmgu> | de Boer et al.,<br>2003 <sup>36</sup> |

We generated a new GM-CSF KO mouse. Specifically, we used the ‘knockout-first’ allele (Csf2tm2a(EUCOMM)Hmgu, MGI: 5497329) design to be able to produce reporter knockouts, conditional knockouts, and null alleles following exposure to site-specific recombinases Cre and Flp.

Mice were generated at LTK Zürich (2018) using genetically modified JM8A3.N1 C57BL/6 stem cells from EUCOMM. ES cells were injected into C57BL/6 blastocysts and the resulting chimeric mice were bred with C57BL/6 to achieve germ line transmission and generate the knockout-first allele. The L1L2_GT0_LF2A_LacZ_BetactP_neo cassette was inserted by targeted mutation and subsequently the knockout-first mice were crossed with deleter1-cre to create the *Csf2*-LacZ strain. The *Csf2*^fl/fl^ mice were created by crossing to the flp delete cre resulting in a conditional ready allele.

*Csf2*-LacZ mice have a disrupted GM-CSF expression and express the LacZ reporter cassette instead. This allows us to simultaneously abrogate GM-CSF expression and analyze the affected cells. *Csf2*^fl/fl^ mice allow us to conditionally abrogate GM-CSF expression in individual cell types by using cell type specific Cre strains.

All mice were kept in a specific pathogen free (SPF) animal facility led by LASC at the University of Zurich. All procedures were approved by the Cantonal Veterinary office and performed under the licenses 159/18 and ZH055/2022. Mice of both sexes were used for all experimental timepoints. If essential age and sex of the mice is noted for the according experiment in the relevant figure legend. Transgenic mice used in this study are listed in table 1.

### Tissue preparation for flow cytometry

Mice were euthanized with CO2 asphyxation, blood was collected from the heart with insulin syringes and following the mice were transcardially perfused with PBS. Red blood cells were removed from the blood with ACK lysis buffer (buffer details). The submandibular and sublingual salivary gland (SG), lacrimal gland (LG) and mammary gland (MG) were harvested after removal of the associated lymphnodes. The tissue was cut into small pieces, transferred to a 6-well plate and digested with 0.4 mg/ml Collagenase IV (#9001-12-1, Sigma-Aldrich) and 0.2 mg/ml deoxyribonuclease I (DNase I) (#E1010, Luzerna) in HBSS (with Ca2^+^ and Mg2^+^) (#14025-050, Gibco) + 10% FCS for 20 min at 37°C. To support, the tissue was mechanically dissociated using a 19-gauge needle and digested for another 20min at 37°C. The tissue was homogenized with a 19-gauge needle and filtered through a 100 μm cell strainer (800100, Bioswisstec) into a 50mL Falcon tube. Washed with cold PBS and fill up to 30mL. Centrifuge at 4°C, 1500rpm for 10min, discard supernatant and resuspend pellet in 200μl and take everything for staining. For BM isolation, BM cells in the femur were flushed out with a 10-ml syringe with PBS through a 100um cell strainer and washed with 30ml PBS. Centrifuge at 4°C, 400xg for 10min, discard supernatant and resuspend pellet in 200μl and take everything for staining.

#### Collagenase Solution

HBSS (with Ca++), 10% FCS, Collagenase type 4 (0.4 mg/ml Collagenase)(Sigma C5138)+ DNAse I (0.2mg/ml)

### Spectral Flow Cytometry

Prior to surface labelling, cells were incubated with a purified anti-mouse CD16/32 (FcBlock, clone 93, Biolegend) for 10 min on ice to prevent non-specific binding of primary antibodies. Single-cell suspensions were then directly incubated with the primary surface antibody cocktail in PBS for 25 min at 4°C. Following a wash step with PBS (1500rpm, 5min, at 4°C) the cells were then incubated in the secondary surface antibody cocktail (fluorochrome-conjugated streptavidin) for 20 min at 4°C. Doublet discrimination was performed based on SSC-A/H, FSC-A/H and dead cell exclusion by using a Fixable Viability Kit (Zombie Nir™, Biolegend, Cat# 423105, dilution 1:400).

Data were acquired on a 5L Aurora spectral analyser (Cytek Biosciences). Flourescence-activated cell sorting was performed using a BD FACS Symphony S6 5L. Flow cytometry data were analyzed using the FlowJo software (version 10.8.0, Tree Star Inc.) and Rstudio (version 4.0.1).

Following anti-mouse fluorochrome-conjugated monoclonal antibodies (mAbs) were used in this study

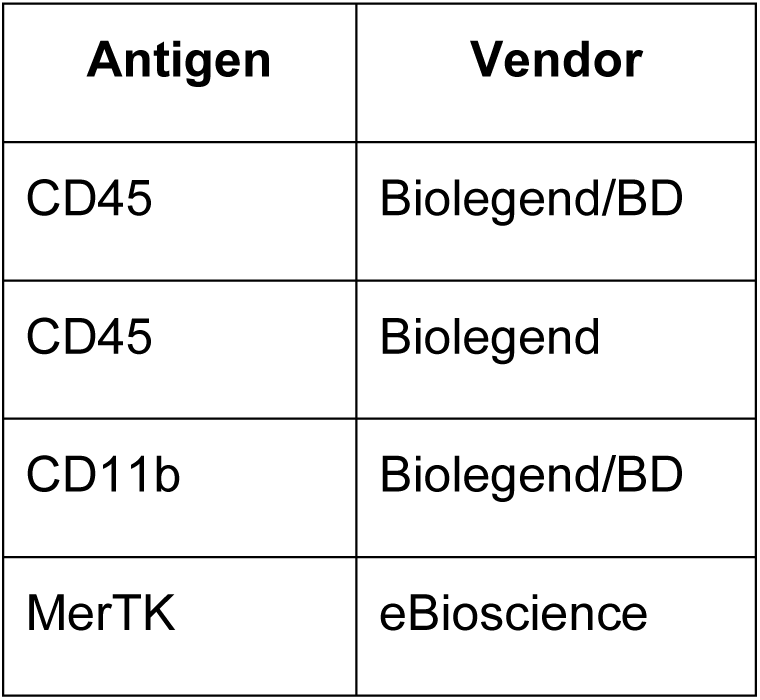
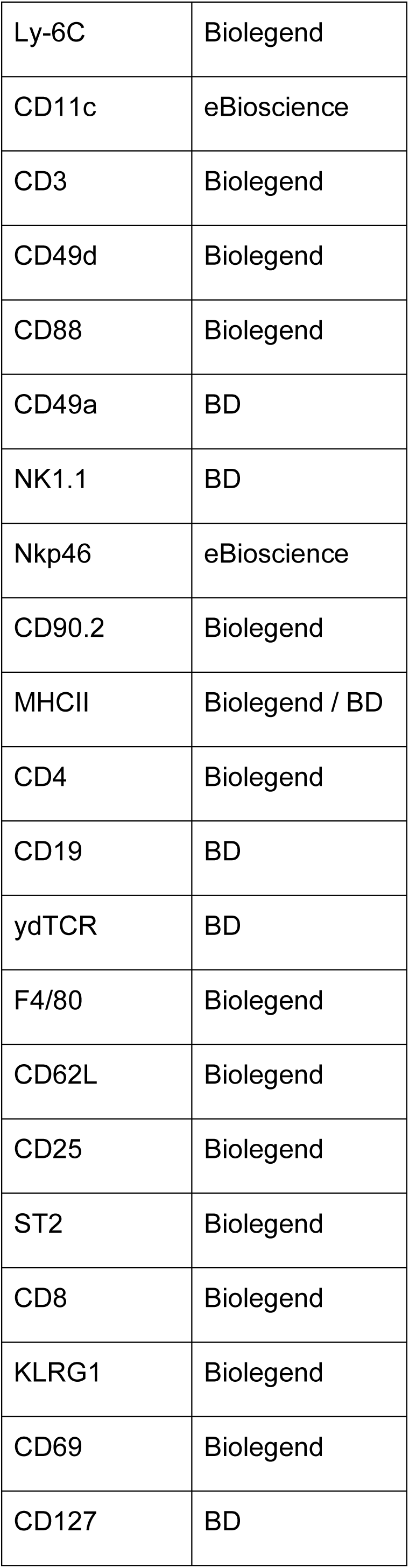
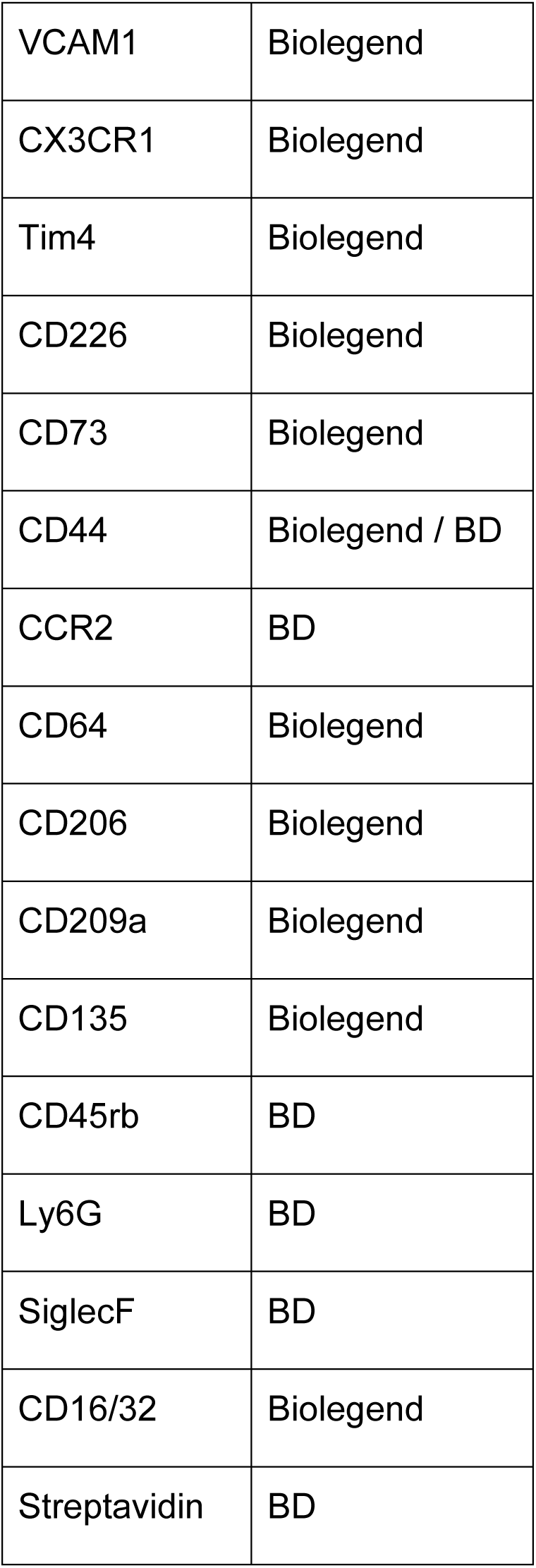

| Antigen | Vendor |
| --- | --- |
| CD45 | Biolegend/BD |
| CD45 | Biolegend |
| CD11b | Biolegend/BD |
| MerTK | eBioscience |
| Ly-6C | Biolegend |
| CD11c | eBioscience |
| CD3 | Biolegend |
| CD49d | Biolegend |
| CD88 | Biolegend |
| CD49a | BD |
| NK1.1 | BD |
| Nkp46 | eBioscience |
| CD90.2 | Biolegend |
| MHCII | Biolegend / BD |
| CD4 | Biolegend |
| CD19 | BD |
| γδTCR | BD |
| F4/80 | Biolegend |
| CD62L | Biolegend |
| CD25 | Biolegend |
| ST2 | Biolegend |
| CD8 | Biolegend |
| KLRG1 | Biolegend |
| CD69 | Biolegend |
| CD127 | BD |
| VCAM1 | Biolegend |
| CX3CR1 | Biolegend |
| Tim4 | Biolegend |
| CD226 | Biolegend |
| CD73 | Biolegend |
| CD44 | Biolegend / BD |
| CCR2 | BD |
| CD64 | Biolegend |
| CD206 | Biolegend |
| CD209a | Biolegend |
| CD135 | Biolegend |
| CD45rb | BD |
| Ly6G | BD |
| SiglecF | BD |
| CD16/32 | Biolegend |
| Streptavidin | BD |

### Tissue preparation for bulkRNAsequencing of myoepithelial cells

The SMG and SLG (SG) were processed according to the previously described protocol with slight adaptations^72^. Minced SG were transferred into a 50 mL conical tube containing digestion medium (2.5mL; DMEM/F12HAM with collagenase type I (750 U/mL), hyaluronidase (500 U/mL), and DNase I (0.1 mg/mL)). Pipette the tissues up and down every 10 min with a 1000ul pipette. After 45 min at 37 °C (shaking at 100/min) centrifuge at 100 × *g* for 20 sec at 4 °C. Aspirate the supernatant and wash twice with 10 mL of DMEM/F12HAM and repeat centrifugation at 100 x g for 20sec at 4°C. After washing the cells, add 5 mL of TrypLE Express and 50 μL of Dnase I(10 mg/mL), carefully pipette with P200, and dissociate for 10 min at 37 °C with shaking (at 100/min speed). Add 50 μL DNase I (10 mg/mL) every 2 min. Centrifuge at 300 *× g* for 5 min at 4 °C. Aspirate the supernatant and wash twice with 10 mL of DMEM/F12HAM and centrifuge at 300 *× g* for 5 min at 4 °C. After washing the cells, add 1 mL of Dispase (2.6 U/mL), 20 μL of DNase I (10 mg/mL), and 9 mL of DMEM/F12HAM (10% FBS) to the tube and carefully pipette with P200 for 1 min. Centrifuge at 350 *× g* for 5 min at 4 °C. Aspirate the supernatant and wash with 10 mL DMEM/F12HAM containing 10% FBS, before centrifugation at 350 *× g* for 5 min at 4 °C. Aspirate the supernatant and resuspend cells in 10 mL DMEM/F12HAM containing 10% FBS. Filter through a 70 μm filter into a new 50 mL tube and then filter through a 40 μm cell strainer into a new 50-mL tube. All cells were stained with fluorescent-conjugated antibodies (anti-TER-119, anti-CD31, anti-CD45, anti-EpCAM, and anti-CD49f antibodies) for 30 min on ice.

The myoepithelial cell fraction (EpCAM^low^ CD49f^high^) was sorted on the BD FACS Symphony S6 5L after excluding the dead cells by using a Fixable Viability Kit (Zombie Nir™, Biolegend, Cat# 423105, dilution 1:400) and the stromal cell populations (TER-119^−^CD31^−^CD45^−^). Following, the RNA was extracted from the sorted MECs using the Qiagen RNeasy Plus Micro Kit (Ref# 74034).

### Bulk RNA Sequencing and analysis

Libraries were prepared using the PicoMammalian v3 kit (Takara Bio) and sequenced using an illumina NovaSeq X plus with 150-bp paired-end reads.

Transcript abundances were quantified via Kallisto 0.50.0^73^ in paired-end mode. Transcript abundances were loaded into R v4.4.1. via tximport v1.32.0^74^ and summarized to gene level via the function summarizeToGene using a matching table of ensemble transcript ids with gene names generated via biomaRt v2.60.1.^75^ Genes with TPM < 2 in 4 or more samples were excluded from further analysis. Differentially expressed genes were calculated with edgeR v4.2.0^76^ by employing a general linear model via the functions glmFit and glmLRT. Genes with FDR adjusted p-values < 0.05 were considered significant. Gene scores were calculated using the AddGeneSetScore function^77^. Heatmaps were generated using the pheatmap v1.0.12 (CRAN).

### Tamoxifen treatment

Tamoxifen was reconstituted with 100% ethanol in corn oil with a final concentration of 25 mg ml–1. A total of 5 mg of tamoxifen was administered via oral gavage to pregnant mice.

### Wright-Giemsa staining

Sort-purified GM-MACS, MHCII+MACS and DCs from P20 SG were spun onto glass slides using a Cytospin 4 Cytocentrifuge (Thermo Scientific), then air-dried at room temperature for 20 min. Slides were stained with Wright-Giemsa solution A for 1 min and Wright-Giemsa solution B for 5 min, then washed with flow water for 5 min. Slides were air-dried and sealed with mounting medium. Images were acquired with an Olympus BX53 light microscope equipped with a 100× oil immersion objective lens. Wright-Giemsa staining was performed by the VetSuisse Pathology, UZH Zürich.

### SEM

Sorted cells were fixed with 4% formaldehyde (28906, Thermo Scientific, Rockford, USA) in 0.1 M sodium cacodylate buffer (pH 7.35, C0250, Sigma-Aldrich, Buchs, Switzerland). Subsequently, cells were fixed with 2.5% glutaraldehyde (G5882, Sigma-Aldrich, Buchs, Switzerland) in 0.1 M sodium cacodylate buffer. 80uL of the solution containing the cells were centrifuged on 12 mm cover glasses using a cytospin centrifuge (6 min at 40 g, Cytospin 2, Thermo Fisher Scientific, Schlieren, Switzerland). Subsequently, samples were rinsed 3 times with PBS, incubated in 1% osmium tetroxide (OsO4, 19152, EMS, Hatfield, USA) in PBS for 30 minutes (OsO4/PBS 1:2), rinsed 3 times with PBS, dehydrated in 70% ethanol (20821.321, VWR Chemical) in H2O for 1 h, 100% ethanol for 1 hour, hexamethyldisilazane (440191, Sigma-Aldrich, Buchs, Switzerland) for 1 hour, and transferred to fresh hexamethyldisilazane prior to air-drying overnight. Dry samples were mounted on aluminium stubs with double sided conductive tape (TAMS, Schwerzenbach, Switzerland) and sputter coated with 4 nm of platinum (working distance 5 cm, argon pressure 1.0e-1 mbar, sputter current 30 mA, CCU-010 high-vacuum coater, Safematic, Zizers, Switzerland) and imaged in a Zeiss GeminiSEM 450 scanning electron microscope (Zeiss, Oberkochen, Germany) at 5 kV acceleration voltage using the secondary electron detector.

### Tissue processing for Histology

Mice were euthanized through CO2 asphyxiation and transcardially perfused with ice-cold PBS. SGs were carefully removed, fixed in 4% PFA (# 11762.00500, Morphisto) for 1-6 hours at 4°C, rinsed in PBS and then incubated in 30% sucrose in PBS at 4°C overnight.

### Immunofluorescence staining and analysis

Salivary or lacrimal glands were embedded in OCT compound (Leica). 10 mm sections were cut using a cryostat (Leica) and stored at −20 °C. Tissue was blocked for 2 hours at room temperature with 5% BSA (Sigma Aldrich), 5% Donkey Serum (Merck) in 0.01% PBS-Tween-20 (PBST). Sections were incubated with primary antibodies (see below) overnight at 4°C. Antibodies were detected using donkey Cy2-, Cy3 or Cy5-conjugated secondary Fab fragment antibodies (Jackson Laboratories) at room temperature for 2 hours. Where applicable, sections were incubated with directly-conjugated primary antibodies overnight at 4°C. Nuclei stained using Hoechst 33342 (1:1000, Sigma Aldrich) for 15 minutes, and slides were mounted using Prolong Gold anti-fade mounting media. Images were acquired on a Leica SP8 confocal microscope and subsequently analysed using NIH ImageJ software.

Following antibodies were used for immunofluorescent staining

| Antibody | Species | Supplier |
| --- | --- | --- |
| αSMA | Mouse | Sigma Aldrich |
| AQP5 | Rabbit | Abcam |
| CD226 | Rat | BioLegend |
| IBA1 | Rabbit | Antibodies Online |
| KRT8 | Rabbit | Sigma Aldrich |
| I-A/I-E 488 (MHCII) | Mouse | BioLegend |
| PDGFRα | Goat | R&D Systems |

### Human salivary gland analysis

Human submandibular gland biopsies were collected with patient consent during neck dissection surgery, from tissue which would otherwise be discarded, under NHS Lothian Tissue Governance approval number SR-857 (Emmerson). Tissue was fixed for 6 hours in 4% paraformaldehyde solution (Sigma Aldrich). Biopsies were further processed as described above for mouse samples.

Following antibodies were used for immunofluorescent staining

| Antibody | Species | Supplier |
| --- | --- | --- |
| αSMA | Mouse | Sigma Aldrich |
| CD226 | Rat | BioLegend |
| IBA1 | Rabbit | Antibodies Online |

### CODEX staining

5µm slices of frozen SG from P14 and P20 FROG mice were prepared and used for CODEX staining as described before (PMID: 36623939). Briefly, sections were dried on drierite beads for 15 min, and fixed for 10 min in ice-cold acetone. Samples were subsequently rehydrated in hydration buffer and photobleached. After photobleaching, sections were blocked and stained with a 30-plex CODEX antibody panel (see below) overnight at 4°C. After staining, samples were washed with staining buffer, fixed with ice-cold methanol, washed with 1x PBS and fixed with BS3 fixative (Sigma Aldrich, St. Louis, MO, USA, Cat# 21580). Samples were washed with 1x PBS and stored at 4°C before imaging.

Following reagents were used to prepare samples for CODEX staining

| Reagent | Producer | Identifier |
| --- | --- | --- |
| CODEX staining kit | Akoya Biosciences | Cat# 7000008 |
| CODEX conjugation kit | Akoya Biosciences | Cat# 7000009 |
| CODEX Buffer | Akoya Biosciences | Cat# 7000001 |
| Nuclear staining | Akoya Biosciences | Cat# 7000003 |
| BS3 fixative | Thermo Fisher Scientific | Cat# 21580 |
| Acetone | Sigma Aldrich | Cat# 650501-1L |
| Methanol Free Formaldehyde | Sigma Aldrich | Cat# 28908 |
| Methanol for HPLC >99.9% | Sigma Aldrich | Cat# 34860-1L-R |

Following antibodies were used for the CODEX panel

| Antibody | Producer |
| --- | --- |
| Rat anti-mouse CD64 | Biolegend |
| Rat anti-mouse CD31 | Biolegend |
| Rat anti-mouse CD45 | Biolegend |
| Rat anti-mouse CX3CR1 | Biolegend |
| Rat anti-mouse F4/80 | Thermo Fisher Scientific |
| Rat anti-mouse Ly6C | Biolegend |
| Mouse anti-mouse SMA | Biolegend |
| Rat anti-mouse CD8a | Biolegend |
| Rat anti-mouse EpCAM | Biolegend |
| Hamster anti-mouse CD11c | Biolegend |
| Rat anti-mouse TCRβ | Biolegend |
| Rat anti-mouse Ly6G | Biolegend |
| Rat anti-mouse CD206 | BioRad |
| Rat anti-mouse CD169 | Biolegend |
| Rat anti-I-A/I-E | Biolegend |
| Rat anti-mouse CD11b | Biolegend |
| Rat anti-mouse CD172a | Biolegend |
| Rat anti-mouse CD90.2 | Biolegend |
| Rat anti-mouse PDGFRa | Thermo Fisher Scientific |
| Rat anti-mouse CD4 | Biolegend |
| Rat anti-mouse CD19 | Biolegend |
| Rat anti-mouse Xcr1 | Biolegend |
| Rat anti-mouse CD26 | Biolegend |
| Rat anti-mouse Siglec F | Thermo Fisher Scientific |
| Rat anti-mouse ST2 | Biolegend |
| Rat anti-mouse CD226 | Biolegend |
| Mouse anti-mouse NK1.1 | Biolegend |
| Rabbit anti-GFP | Thermo Fisher Scientific |
| Rabbi anti-RFP | Rockland |

### CODEX imaging, processing and analysis

Coverslips were equilibrated at room temperature before imaging. A multicycle CODEX experiment was performed following manufacturer’s instructions (Akoya Biosciences). Images were acquired with a Zeiss Axio Observer widefield fluorescence microscope using a 20x objective (NA 0.85). A total of 6 slices with a z-spacing of 1.5 µm were acquired. The 405, 488, 568, and 647 nm channels were used. Raw files were exported using the CODEX Instrument Manager (Akoya Biosciences, Marlborough, MA, USA) and processed with CODEX Processor v1.7 (Akoya Biosciences).

Images were analyzed in the CODEX MAV software (Akoya Biosciences). First, artefacts identified as objects were removed using DAPI signals as a reference. Afterwards, a manual cell classification was performed using the same gating strategy as the one used in flow cytometry. After cell classification, a LogOddRatio analysis to determine the probability of interaction between cell types was performed. The minimum and maximum distances of interaction were 5 and 30 µm, respectively. A similar procedure was performed to determine the interaction of adenophages with tdTomato^+^ lymphoid cells. To validate the latter, images were visualized in QuPath. A composite classifier for adenophages was generated and used to detect these cells in the image. tdTomato^+^cells neighboring adenophages were inspected and identified based on their marker expression.

### Single-cell RNA-sequencing (10× Genomics)

For scRNA-seq of P20 SG, tissue was digested as described above. Total CD45+ cells were pre-selected using CD45 MicroBeads (Milteny Biotec, #130-052-301) on LS columns and following further sort-purified for CD45+ cells. Each sample was pooled from two individual P20 mice.

Total CD45+ cells were pelleted and resuspended in PBS containing 0.04% BSA at a final density of ∼ 1,000 cells per μl, these cell suspensions were subjected for scRNA-seq analysis using 10X× Genomics Chromium Single Cell 3’ Reagent Kit (v3) according to the manufacturer’s instructions. Libraries were sequenced using an illumina Hiseq x Ten or Novaseq 6000 sequencer with 150-bp paired-end reads. The sequencing data were processed with Cell Ranger Single Cell Software Suite provided by 10X× Genomics. The output matrics were further analysed in R mostly with the Seurat package^78^. Lowly-expressed genes (detected in fewer than 3 cells) were filtered out, cells with less than 800 nfeatures or with high mitochondrial gene counts s (>15% of counts per cell) were removed.

To compare the gene expression profiles across different clusters from multiple datasets (scRNA-seq comparisons from different datasets), we first calculated the mean expression value in each cluster. Then we carried out quantile normalization on the tpm matrix to make data distributions identical, using normalize.quantiles function in R preprocessCore package (version 1.48.0). The normalized data were then processed with ComBat to remove batch effect across different datasets. pvclust function from pvclust package (versio 2.2.0) were used to calculate p-values for hierarchical clustering with “ward.D” method using “euclidean” distance. In Figure 7, we selected top 1000 genes with the highest stand deviation in the batch-removed normalized expression matrix and then calculated the Pearson correlation across clusters from human and mouse scRNA-seq datasets. To generate the heatmap, we z-transformed the correlation matrix by column.

For selection of clustering parameters we iterated over several combinations of clustering resolution and PCA components used for dimensional reduction on the basis of the R package Harmony normalization^79^. We selected a clustering resolution of 0.65 and 15 PCA components based on the infliction point of the elbow plot generated using Seurat.

For projection of our dataset onto the Tabular Muris dataset^80^ we employed the mapQuery function from the R package Symphony^81^ as well as the knnPredict.Seurat from Seurat. Proportions barplots were generated with the R package Platypus^82^.

For integration of published single cell RNA-seq datasets with our single cell RNA-seq data, we use the R package STACAS^83^ with 2000 anchor.features and 15 PCA dimensions. To include all genes present in both dataset in the final integrated object, we provided the overlapping genes as features.to.integrate in the Integrated.Data.STACAS utility.

CellphoneDB^51^ analysis was run in statistical_analysis mode with a threshold setting of 0.1.

All plots were generated using the R package ggplot2 (Hadley Wickham, 2016). Color palettes were from the R packages Pals (Kevin Wright, 2023) and MetColorBrewer (Blake Robert Mills, 2022).

### Single nuclei preparation and Fluorescence activated nuclei sorting

Salivary gland was harvested from P20 pups and immediately snap-frozen in liquid nitrogen and stored at −80°C until further processing. Nuclei were isolated with the EZ PREP lysis buffer (#NUC-101, Sigma). Tissue samples were homogenized using a glass dounce tissue grinder (#D8938, Sigma) (25 strokes with pastel A, 25 strokes with pastel B) in 2 ml of ice-cold EZ prep lysis buffer and incubated on ice with an additional 2 ml (we added 1 ml for SGϑ) of ice-cold EZ PREP lysis buffer. During incubation, 1 μΜ of Hoechst (#H3570, Thermo Fisher Scientific) dye and 40 U/μl of RiboLock inhibitors (#EO0382, Thermo Fisher Scientific) were added to the homogenate. Following incubation, the homogenate was filtered through a 30 μM FACS tube filter. Nuclei were sorted based on the fluorescent Hoechst signal using a BD FACS Aria sorter III with an 85 μm nozzle configuration at 4°C, directly into PBS + 4% BSA + RiboLock inhibitors (40 U/μl) (Cat # EO0382). As CNS nuclei vary strongly in size, no doublet exclusion was performed based on FSC or SSC to avoid bias against nucleus size. Nuclei were then counted based on brightfield image and Trypan Blue staining (Cat # 15250061) using a Neubauer counting chamber and a Keyence BZX-710 microscope.

### Droplet based single-nucleus RNA sequencing

Immediately after extraction, 17.500 sorted nuclei per sample were loaded onto a Chromium Single Cell 3′ Chip (10X Genomics) and processed for the single-nucleus cDNA library preparation (Chromium Next GEM Single Cell 3’ Reagent Kits v3.1 protocol). 50.000 reads per nucleus were sequenced using the Illumina Novaseq 6000 #1 platform according to the manufacturer’s instructions without modifications (R1= 28, i7= 10, i5= 10, R2=90). Preparation of cDNA libraries and sequencing were performed at the Functional Genomics Center Zurich. CellRanger software (v6.0.2) was implemented for library demultiplexing, barcode processing, fastq file generation, gene alignment to the mouse genome (GENCODE reference build GRCh38.p6 Release M23), and unique molecular identifier (UMI) counts. We implemented the “include-introns” option for counting intronic reads, as the snRNA-seq assay captures unspliced pre-mRNA as well as mature mRNA. For each sample, a CellRanger report was obtained with all the available information regarding sequencing and mapping parameters. All samples were merged into a matrix using CellRanger (cellranger -aggr function).

For integration of single cell and single nuclei RNA-seq datasets, as well as published single cell RNAseq datasets with our single cell RNA-seq data, we use the R package STACAS (Andreatta et al., 2021) with 2000 anchor.features and 15 PCA dimensions. To include all genes present in both dataset in the final integrated object, we provided the overlapping genes as features.to.integrate in the Integrated.Data.STACAS utility.

### Phagocytosis assay

SG cells were isolated as described above. Following isolation, cells were stained as described above, resuspended in RPMI with 2% FBS and incubated with pHrodo Red *S. aureus* BioParticles (Invitrogen) (0.01 mg ml^−1^, 0.05 mg ml^−1^ and 0.01 mg ml^−1^, respectively), for 1.5 h at 37 °C and 5% CO2. After the incubation, cells were immediately acquired.

### In vitro T cell proliferation assays

A total of 1×105 sort-purified naive TCRβ+CD4+ OT-II T cells were labeled with CellTrace Violet according to the manufacturer’s instructions and co-cultured with sort-purified GM-MACS, MHCII+MACS, DCs or Bcells. The APCs were pulsed with OVA323–339 (10 μg/ml) for 1h at 37°C. Dilution of the cell dye was assessed by flow cytometry on day 3 of culture.

---

**Supplementary Figure 1:**
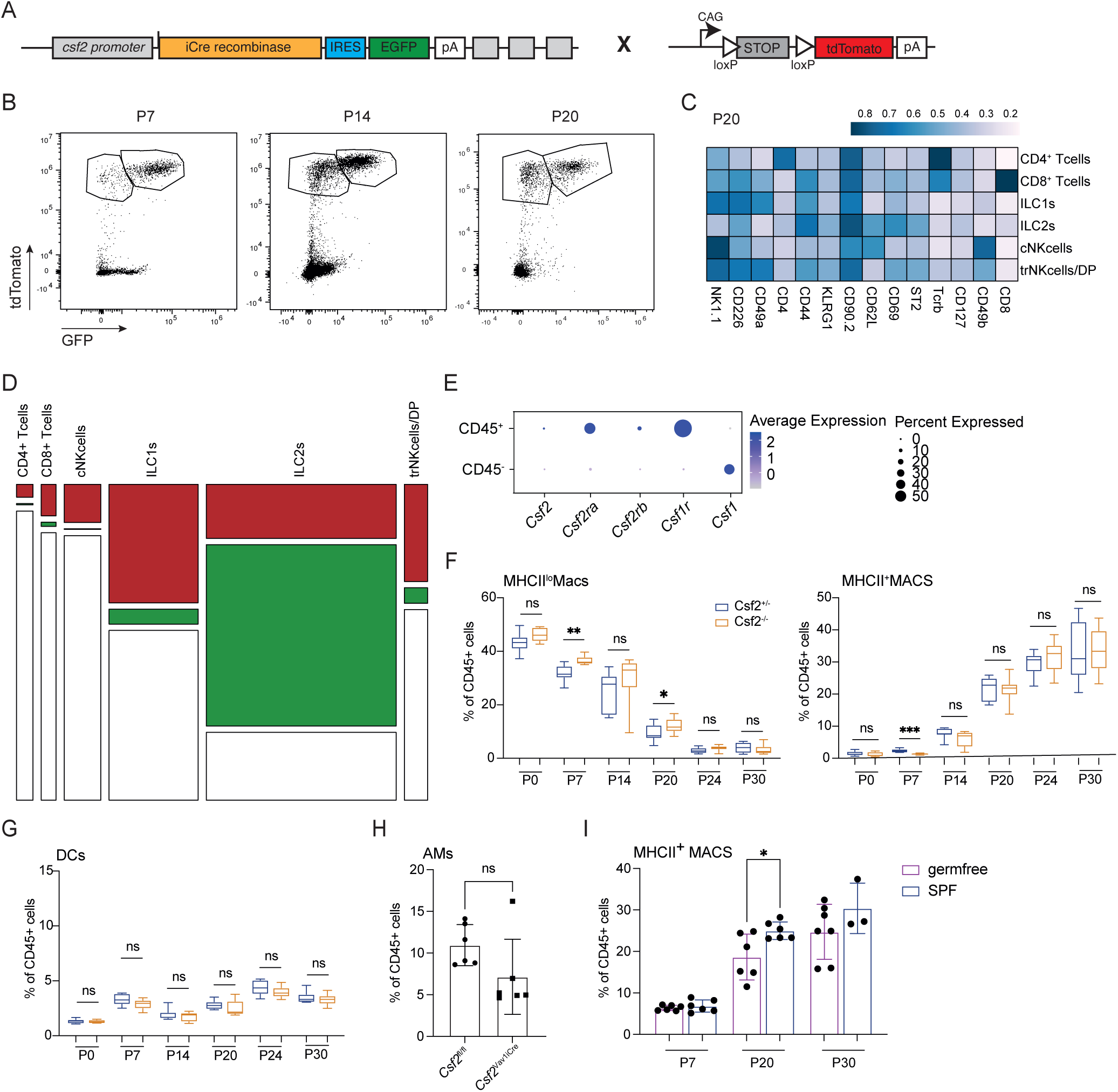
Lymphocytes are the exclusive source of GM-CSF for adenophages. A) Genetic construct of the FROGxAi14 mouse model. B) Representative FACS plots for tdTomato^+^ and tdTomato^+^GFP^+^ cells in the SG of FROGxAi14 mice at P7, P14 and P20. C) Marker expressions in lymphoid populations identified in the SG of FROGxAi14 mice at P20 D) Mosaic plot showing the fraction of tdTomato^+^ (red; ex-GM-CSF producing) and tdTomato^+^GFP^+^ (green; GM-CSF producing) cells in the different lymphocyte populations in the SG of FROGxAi14 mice at P20. The width of the columns represent the abundance of the indicated populations. Gated on liveCD45^+^CD11b^−^. E) scRNAseq and snRNaseq data showing gene expression of indicated genes in the SG of Csf2^+/−^ mice at P20 SG. Colors indicate the average expression of each gene, circle sizes represent percentage of cells within throughout postnatal development in the SG of *Csf2*^+/−^ (blue) and *Csf2*^−/−^ (orange) mice. Pre-gated on live CD45^+^Lin^−^Ly6G^−^SiglecF^−^Ly6C^−^CD11b^+^F4/80^+^ cells. Data are represented as mean±SD. Statistical test: 2-way ANOVA. n = 6-12 mice per group and timepoint from 2-3 independent experiments. G) Frequencies of DCs throughout postnatal development in the SG of *Csf2*^+/−^ (blue) and *Csf2*^−/−^ (orange) mice. Pre-gated on live CD45^+^Lin^−^Ly6G^−^SiglecF^−^Ly6C^−^ F4/80^−^CD11c^+^MHCII^+^ cells. Statistical test: 2-way ANOVA. n = 6-12 mice per group and timepoint from 2-3 independent experiments. H) Frequencies of AMs of *Csf2*^fl/fl^ and *Csf2*^Vav1iCre^ mice at P20. Data are represented as mean±SD. I) Frequency of MHCII^+^ MACS in the SG of germfree (purple) and SPF (blue) mice at indicated time-points. Statistical test: 2-way ANOVA. Data are represented as mean±SD. AM = alveolar macrophages; SPF = specific-pathogen-free.

**Supplementary Figure 2:**
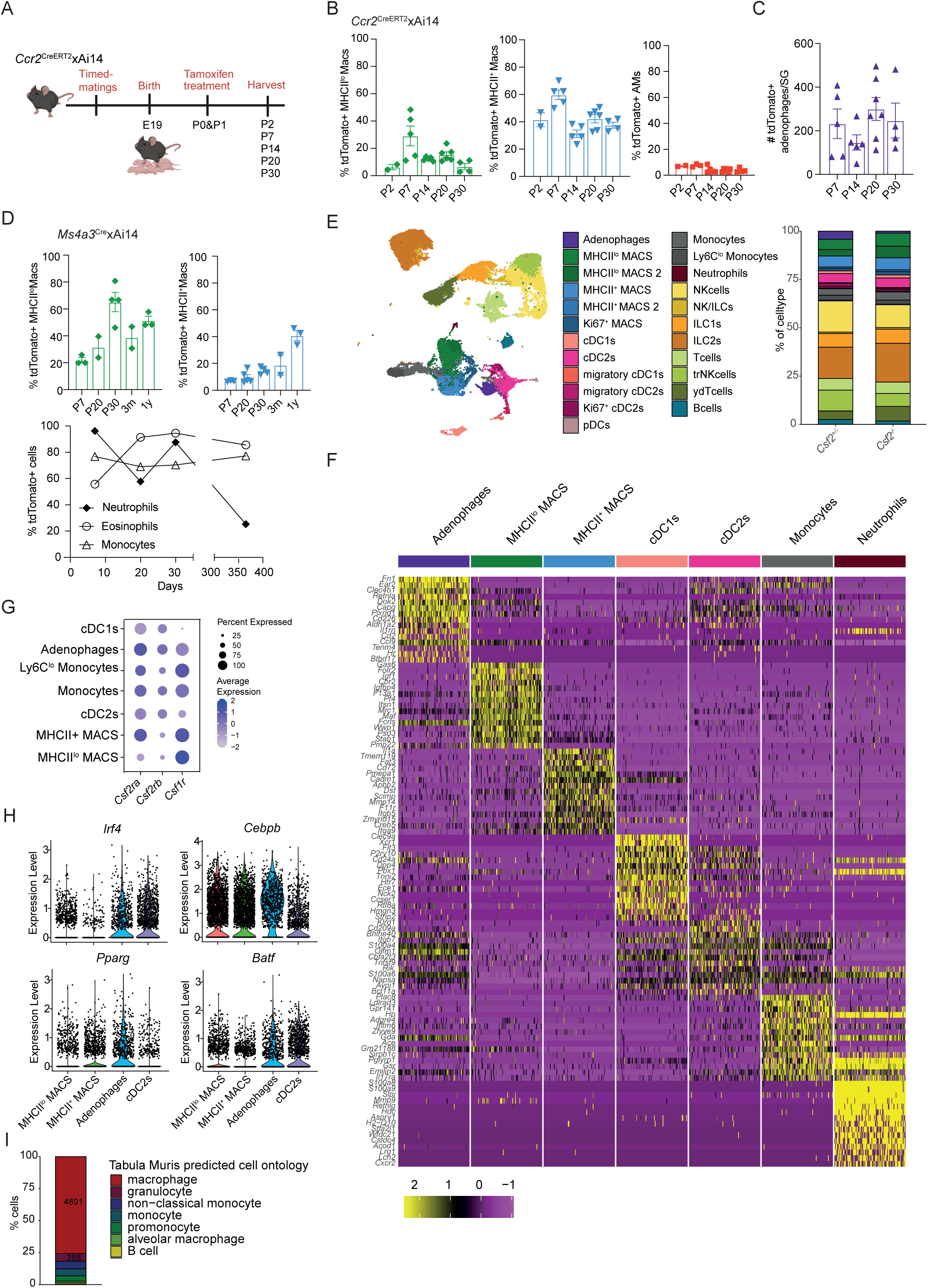
Adenophages arise from fetal monocytes and have a transcriptional signature imprinted by GM-CSF. A) Experimental scheme showing the procedure to label fetal monocyctes in Ccr2^CreERT2^xAi14 mice. B) Frequencies of tdTomato^+^ cells within MHCII^lo^ MACS (green), MHCII^+^MACS (blue) and AM (red) at indicated timepoints after Tamoxifen application at P0/P1 in Ccr2^CreERT2^xAi14 mice. C) Total counts of tdTomato^+^ adenophages in the SG at indicated timepoints after Tamoxifen application at P0/P1 in Ccr2^CreERT2^xAi14 mice. D) top: Frequency of tdTomato^+^ MHCII^lo^ MACS (green) and MHCII^+^ MACS (blue) in the SG of Ms4a3^Cre^xAi14 mice at indicated timepoints. Bottom: Frequency of tdTomato^+^ neutrophils, eosinophils and monocytes in the SG of Ms4a3^Cre^xAi14 mice at indicated timepoints. A-D) Data in bar diagrams are represented as mean±SEM. E) UMAP visualisation of scRNAseq data on all CD45^+^ cells in the SG at P20. Bar diagram visualizes the abundance of the identified immune cell subsets in the SG of Csf2^+/−^ and Csf2^−/−^ mice at P20. F) Gene expression profiles of adenophages, MHCII^lo^ MACS, MHCII^+^ MACS, cDC1s, cDC2s, Monocytes and Neutrophils obtained from scRNAseq data described in E. G) Dot plot showing *Csf2ra*, *Csf2rb* and *Csf1r* expression across indicated myeloid subsets. Colors indicate the average expression of each gene, circle sizes represent percentage of cells within a cluster expressing the indicated gene. H) Violin plots showing expression of *Irf4, Pparg, Cebpb* and *Batf* in macrophage subsets, adenophages and cDC2 in the SG of Csf2^+/−^ mice at P20. I) Bar diagram shows the immune cell populations proposed by the tabula muris data base based on the signature of adenophages. AM = alveolar macrophages.

**Supplementary Figure 3:**
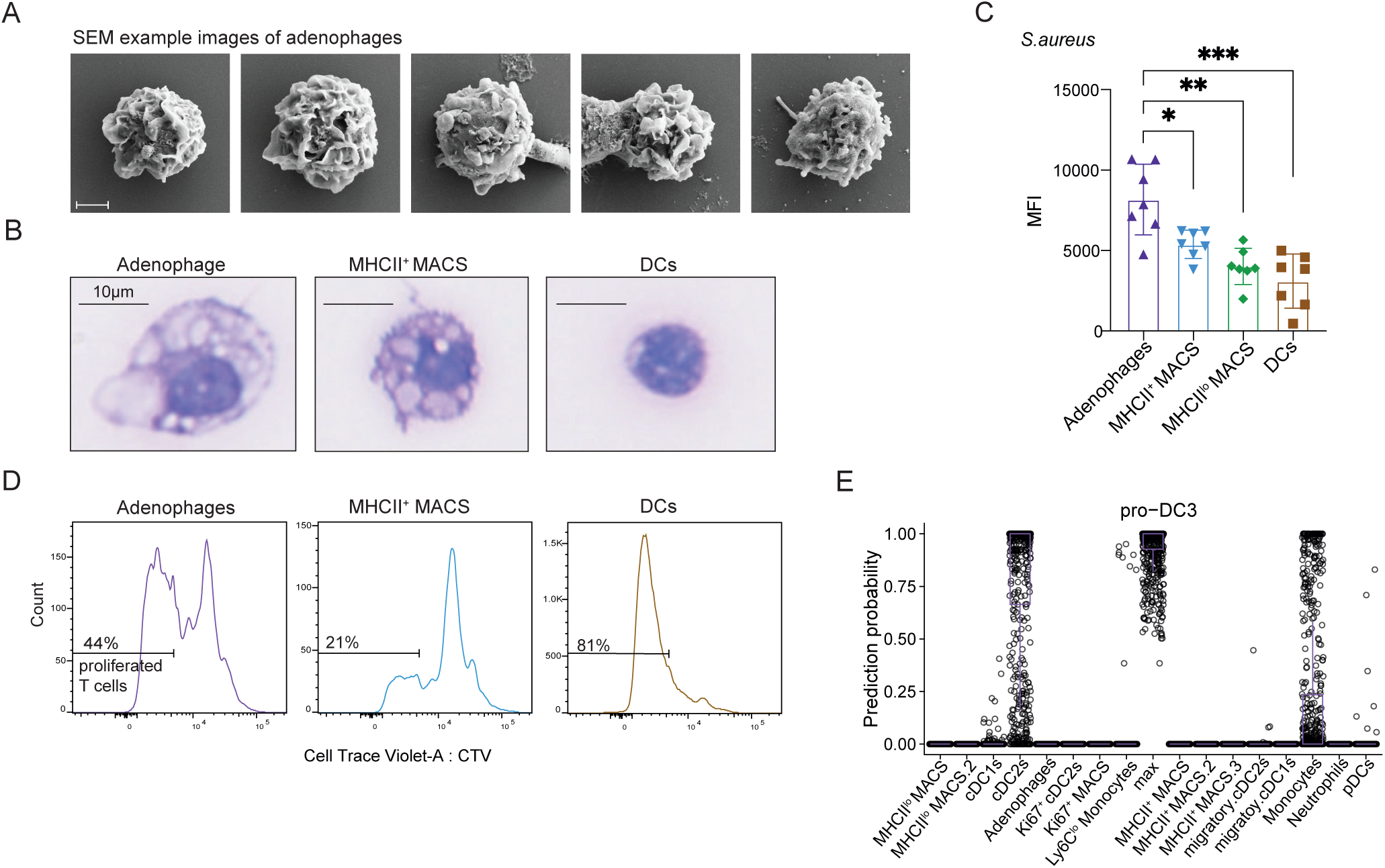
Adenophages share functional and morphological features with DCs and macrophages. A) Additional SEM images of FACS sorted adenophages. Scale = 2 µm. B) Giemsa staining micrographs of sorted adenophages, MHCII^+^ MACS and DCs from the SG of Control mice at P20, representative images taken from two independent experiments. C) MFI of pHrodo Red^+^ adenophages, MHCII^+^ MACS, MHCII^lo^ MACS and DCs from the P20 SG of *Csf2*^+/−^ mice after *in vitro* exposure to S.aureus pHrodo bioparticles for 180min. Data is pooled from 3 independent experiments Statistics: Kruskal-Wallis test with Dunn’s multiple comparisons test. Data in bar diagram is represented as mean±SD. D) CellTraceViolet (CTV) expression in CD4^+^ OT-II transgenic T cells after co-culture with adenophages, MHCII^+^ MACS or DCs isolated from the SG of Csf2+/− mice at P20 and pulsed with OVA_323–339_ for 1 hour. Data is representative for two independent experiments. E) Probability prediction of proDC3s falling into one of the myeloid clusters identified in the SG of control mice at P20. MFI = mean fluorescent intensity.

**Supplementary Figure 4:**
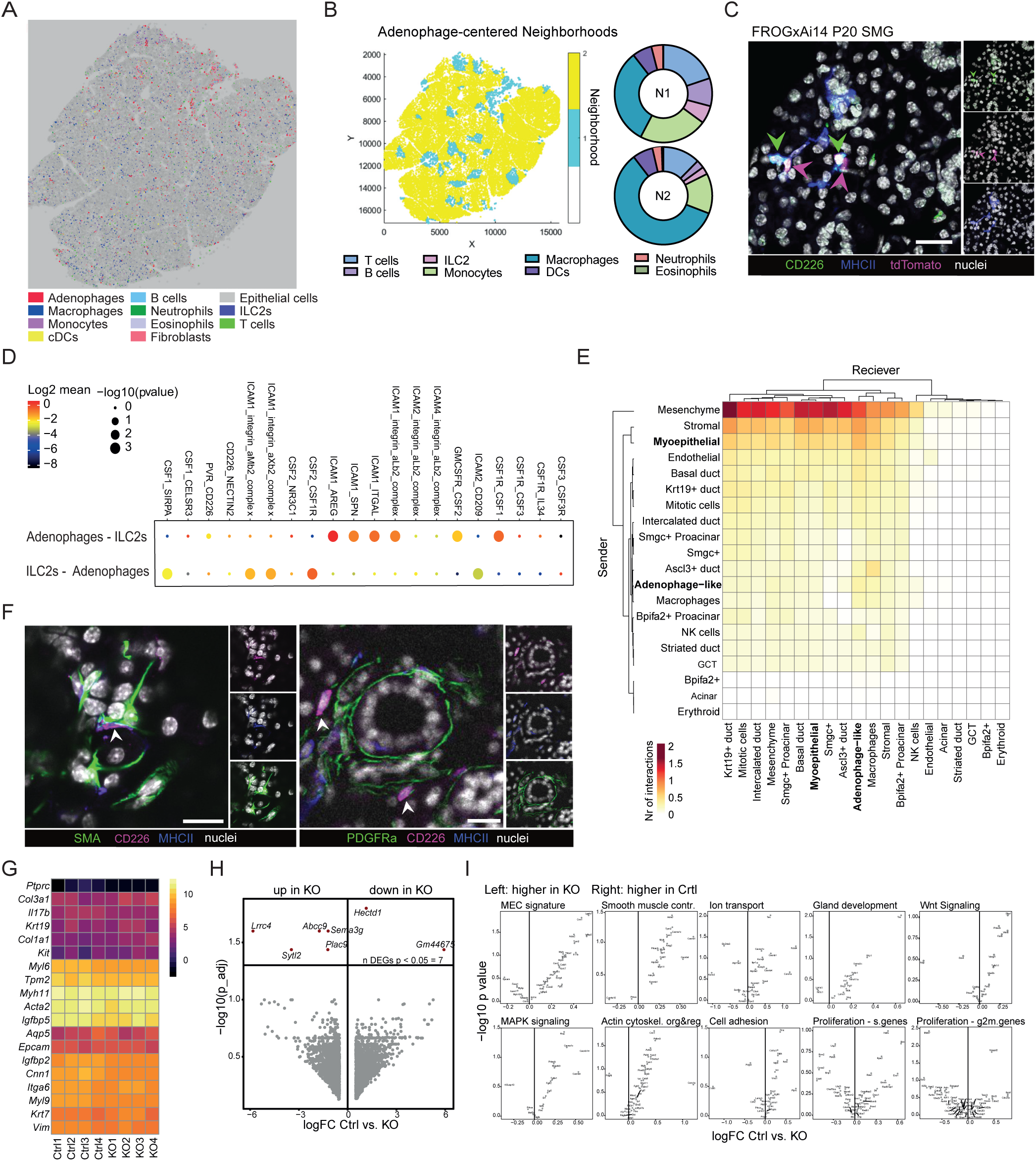
Adenophages have a relationship with ILC2s and MECs. A) CODEX analysis showing the different cellular subsets by indicated color coding. B) Neighborhood analysis of adenophages using the CODEX data. Section heatmap (left) indicates distribution of the neighborhood; pie charts (right) indicate composition of the neighborhoods. C) Immunofluorescent histology image of SMG of FROGxAi14 mice at P20. Green arrow heads indicate adenophages, magenta arrow heads indicate tdTomato^+^ cells (indicating GM-CSF production). Green = CD226, Blue = MHCII, Red = tdTomato, White = DAPI. Scale: 20 µm. D) Cell-Chat analysis of cell-cell interactions between adenophages and ILC2s using scRNAseq data on immune cells in Csf2^+/−^ mice at P20. E) Cell-Chat analysis of the P30 timepoint of the scRNAseq data set from Hauser et al., 2020. Heatmap indicates interaction probability. F) Immunofluorescent histology image of the SMG of wildtype mice at P20. Images are representative for the imaging used to define location of adenophages within the tissue (see Figure 4E). Left image shows staining for αSMA (green); right image shows staining for PDGFRα (green); both images: Magenta = CD226, Blue = MHCII and White = DAPI. Scale: 20 µm. G) Heatmap showing gene expression obtained by RNAseq of the indicated MEC samples Ctrl = Csf2^+/−^; KO = Csf2^−/−^. H) Volcanoplot showing DEGs between MECs isolated from the SG of *Csf2*^+/−^ and *Csf2*^−/−^ mice at P20. I) Volcanoplots show identity and functionality signature genes (see Supplementary Table 1) in MECs isolated from the SG of *Csf2*^+/−^ and *Csf2*^−/−^ mice at P20. SMG = submandibular salivary gland; MEC = myoepithelial cell; DEGs = differentially expressed genes.

**Supplementary Figure 5:**
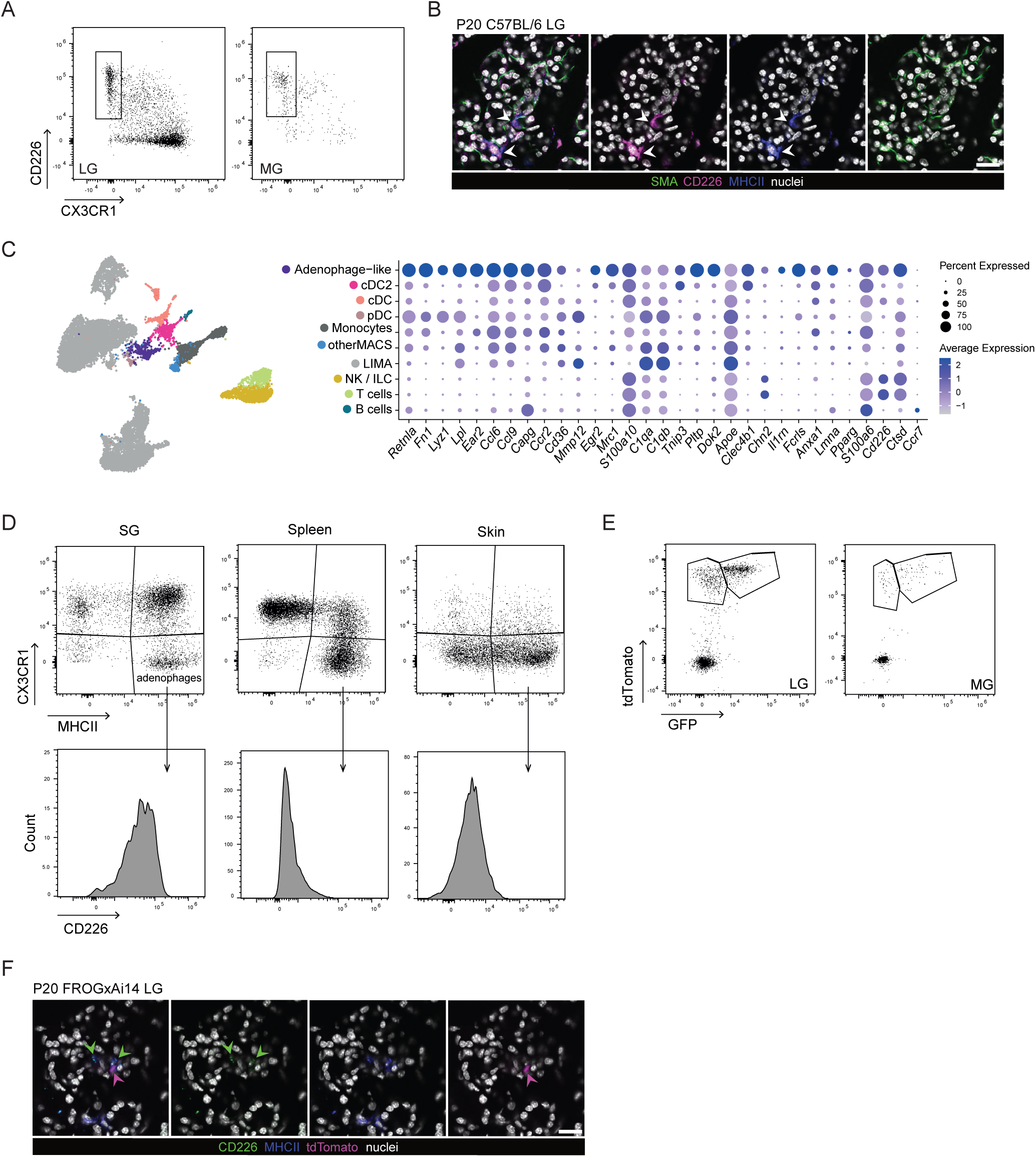
Adenophages are a conserved cell population across glands and species. A) Representative FACS plots for adenophages in the LG (left) and MG (right). Pre-gated on singlets live CD45^+^CD90^−^Ly6G^−^SiglecF^−^Ly6C^−^CD11b^+^F4/80^+^MHCII^+^. B) Immunofluorescent histology image of the P20 LG of wildtype mice. Green = aSMA, Magenta = CD226, Blue = MHCII, White = DAPI. Scale: 20 µm. C) Reanalysis of the scRNAseq dataset on naïve mammary gland (Cansever, Petrova et al., 2023) highlighting the myeloid populations. The dot plot shows the expression of the adenophage signature genes across the myeloid populations. Colors indicate the average expression of each gene, circle sizes represent percentage of cells within a cluster expressing a gene. D) Representative FACS plots of SG macrophages (left), Spleen macrophages (middle) and Skin macrophages (right) with associated histogram for CD226 expression within the CX3CR1^−^MHCII^+^ population. Pre-gated on: live CD45^+^CD90^−^SiglecF^−^Ly6G^−^Ly6C^−^CD11b^+^F4/80^+^CD64^+^. E) Representative FACS plots for tdTomato^+^ and tdTomato^+^GFP^+^ cells in LG (left) and MG (right) of FROGxAi14 mice at P20. Pre-gated on singlet live CD45^+^CD11b^−^ cells. F) Immunofluorescent histology image of the P20 LG of FROGxAi14 mice. Green arrow heads indicate adenophages and magenta arrow heads indicate tdTomato^+^ cells indicating GM-CSF expression. Green = CD226, Blue = MHCII, Red = tdTomato, White = DAPI. Scale: 20 µm. LG = lacrimal gland; MG = mammary gland.

**Supplementary Table 1:**
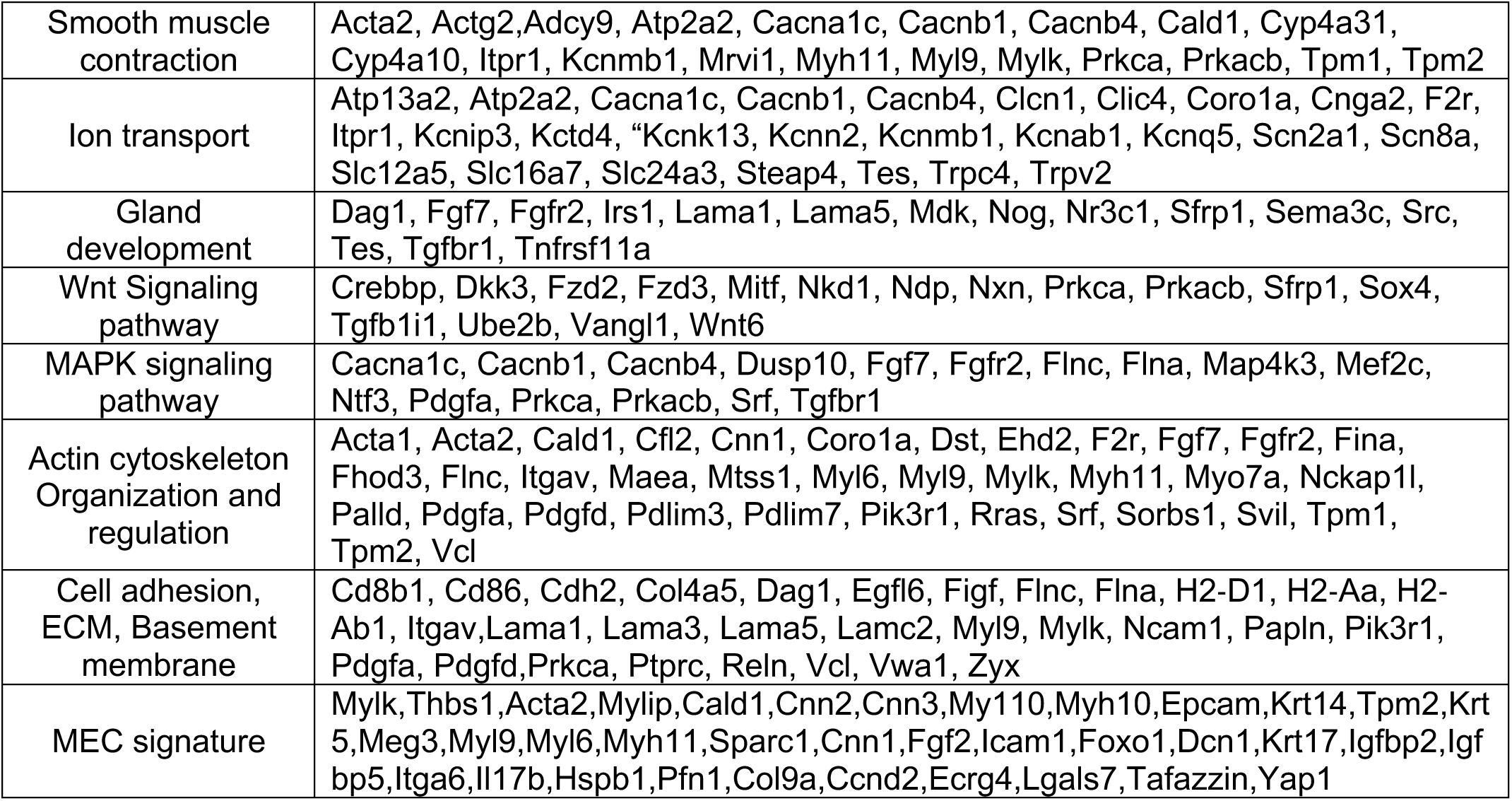
Genes used for MEC Identity Function Scores. List based on: Hauser et al, 2020; Mauduit et al., 2024; Thiemann et al., 2022; Lu et al.,2012

## References

1. Burgess A, Metcalf D. The nature and action of granulocyte-macrophage colony stimulating factors. Blood. 1980;56(6):947–958. doi:10.1182/blood.V56.6.947.947

2. Sallusto F, Lanzavecchia A. Efficient presentation of soluble antigen by cultured human dendritic cells is maintained by granulocyte/macrophage colony-stimulating factor plus interleukin 4 and downregulated by tumor necrosis factor alpha. Journal of Experimental Medicine. 1994;179(4):1109–1118. doi:10.1084/jem.179.4.1109

3. Schneider C, Nobs SP, Kurrer M, Rehrauer H, Thiele C, Kopf M. Induction of the nuclear receptor PPAR-γ by the cytokine GM-CSF is critical for the differentiation of fetal monocytes into alveolar macrophages. Nat Immunol. 2014;15(11):1026–1037. doi:10.1038/ni.3005

4. Gschwend J, Sherman SPM, Ridder F, et al. Alveolar macrophages rely on GM-CSF from alveolar epithelial type 2 cells before and after birth. Journal of Experimental Medicine. 2021;218(10). doi:10.1084/JEM.20210745/212602

5. Guilliams M, De Kleer I, Henri S, et al. Alveolar macrophages develop from fetal monocytes that differentiate into long-lived cells in the first week of life via GM-CSF. J Exp Med. 2013;210(10):1977–1992. doi:10.1084/jem.20131199

6. Bain CC, MacDonald AS. The impact of the lung environment on macrophage development, activation and function: diversity in the face of adversity. Mucosal Immunology 2022 15:2. 2022;15(2):223–234. doi:10.1038/s41385-021-00480-w

7. Greter M, Helft J, Chow A, et al. GM-CSF Controls Nonlymphoid Tissue Dendritic Cell Homeostasis but Is Dispensable for the Differentiation of Inflammatory Dendritic Cells. Immunity. 2012;36(6):1031–1046. 10.1016/j.immuni.2012.03.027

8. King IL, Kroenke MA, Segal BM. GM-CSF–dependent, CD103+ dermal dendritic cells play a critical role in Th effector cell differentiation after subcutaneous immunization. Journal of Experimental Medicine. 2010;207(5):953–961. doi:10.1084/jem.20091844

9. Bogunovic M, Ginhoux F, Helft J, et al. Origin of the Lamina Propria Dendritic Cell Network. Immunity. 2009;31(3):513–525. 10.1016/j.immuni.2009.08.010

10. Chi L, Liu C, Gribonika I, et al. Sexual dimorphism in skin immunity is mediated by an androgen-ILC2-dendritic cell axis. Science (1979). 2024;384(6692):eadk6200. doi:10.1126/science.adk6200

11. Kitamura T, Tanaka N, Watanabe J, Kanegasaki S, Yamada Y, Nakata K. Idiopathic Pulmonary Alveolar Proteinosis as an Autoimmune Disease with Neutralizing Antibody against Granulocyte/Macrophage Colony-Stimulating Factor. Journal of Experimental Medicine. 1999;190(6):875–880. doi:10.1084/jem.190.6.875

12. Uchida K, Nakata K, Trapnell BC, et al. High-affinity autoantibodies specifically eliminate granulocyte-macrophage colony-stimulating factor activity in the lungs of patients with idiopathic pulmonary alveolar proteinosis. Blood. 2004;103(3):1089–1098. doi:10.1182/blood-2003-05-1565

13. Ikegami M, Carter K, Bishop K, et al. Intratracheal Recombinant Surfactant Protein D Prevents Endotoxin Shock in the Newborn Preterm Lamb. Am J Respir Crit Care Med. 2006;173(12):1342–1347. doi:10.1164/rccm.200509-1485OC

14. Berclaz PY, Shibata Y, Whitsett JA, Trapnell BC. GM-CSF, via PU.1, regulates alveolar macrophage FcγR-mediated phagocytosis and the IL-18/IFN-γ–mediated molecular connection between innate and adaptive immunity in the lung. Blood. 2002;100(12):4193–4200. 10.1182/blood-2002-04-1102

15. Cohen M, Giladi A, Gorki AD, et al. Lung Single-Cell Signaling Interaction Map Reveals Basophil Role in Macrophage Imprinting. Cell. 2018;175(4):1031–1044.e18. 10.1016/j.cell.2018.09.009

16. Mortha A, Chudnovskiy A, Hashimoto D, et al. Microbiota-Dependent Crosstalk Between Macrophages and ILC3 Promotes Intestinal Homeostasis. Science (1979). 2014;343(6178):1249288. doi:10.1126/science.1249288

17. Zhao Q, Pan S, Zhang L, et al. A Salivary Gland Resident Macrophage Subset Regulating Radiation Responses. 101177/00220345221150005. Published online March 8, 2023:002203452211500. doi:10.1177/00220345221150005

18. Chiaranunt P, Burrows K, Ngai L, et al. Microbial energy metabolism fuels an intestinal macrophage niche in solitary isolated lymphoid tissues through purinergic signaling. Sci Immunol. 2024;8(86):eabq4573. doi:10.1126/sciimmunol.abq4573

19. Wicks IP, Roberts AW. Targeting GM-CSF in inflammatory diseases. Nat Rev Rheumatol. 2016;12(1):37–48. doi:10.1038/nrrheum.2015.161

20. Panopoulos AD, Watowich SS. Granulocyte colony-stimulating factor: Molecular mechanisms of action during steady state and ‘emergency’ hematopoiesis. Cytokine. 2008;42(3):277–288. 10.1016/j.cyto.2008.03.002

21. Ushach I, Zlotnik A. Biological role of granulocyte macrophage colony-stimulating factor (GM-CSF) and macrophage colony-stimulating factor (M-CSF) on cells of the myeloid lineage. J Leukoc Biol. 2016;100(3):481–489. doi:10.1189/jlb.3RU0316-144R

22. Croxford AL, Lanzinger M, Hartmann FJ, et al. The Cytokine GM-CSF Drives the Inflammatory Signature of CCR2+ Monocytes and Licenses Autoimmunity. Immunity. 2015;43(3):502–514. doi:10.1016/j.immuni.2015.08.010

23. Komuczki J, Tuzlak S, Friebel E, et al. Fate-Mapping of GM-CSF Expression Identifies a Discrete Subset of Inflammation-Driving T Helper Cells Regulated by Cytokines IL-23 and IL-1β. Immunity. 2019;50(5):1289–1304.e6. doi:10.1016/J.IMMUNI.2019.04.006

24. Louis C, Guimaraes F, Yang Y, et al. NK cell–derived GM-CSF potentiates inflammatory arthritis and is negatively regulated by CIS. Journal of Experimental Medicine. 2020;217(5). doi:10.1084/jem.20191421

25. Li R, Rezk A, Miyazaki Y, et al. Proinflammatory GM-CSF–producing B cells in multiple sclerosis and B cell depletion therapy. Sci Transl Med. 2015;7(310):310ra166 LP-310ra166. doi:10.1126/scitranslmed.aab4176

26. Anzai A, Choi JL, He S, et al. The infarcted myocardium solicits GM-CSF for the detrimental oversupply of inflammatory leukocytes. Journal of Experimental Medicine. 2017;214(11):3293–3310. doi:10.1084/jem.20170689

27. Amorim A, De Feo D, Friebel E, et al. IFNγ and GM-CSF control complementary differentiation programs in the monocyte-to-phagocyte transition during neuroinflammation. Nat Immunol. 2022;23(2):217–228. doi:10.1038/s41590-021-01117-7

28. Becher B, Tugues S, Greter M. GM-CSF: From Growth Factor to Central Mediator of Tissue Inflammation. Immunity. 2016;45(5):963–973. doi:10.1016/J.IMMUNI.2016.10.026

29. Codarri L, Gyülvészii G, Tosevski V, et al. RORγ3t drives production of the cytokine GM-CSF in helper T cells, which is essential for the effector phase of autoimmune neuroinflammation. Nat Immunol. 2011;12(6):560–567. doi:10.1038/ni.2027

30. Croxford AL, Spath S, Becher B. GM-CSF in Neuroinflammation: Licensing Myeloid Cells for Tissue Damage. Trends Immunol. 2015;36(10):651–662. doi:10.1016/j.it.2015.08.004

31. Spath S, Komuczki J, Hermann M, et al. Dysregulation of the Cytokine GM-CSF Induces Spontaneous Phagocyte Invasion and Immunopathology in the Central Nervous System. Immunity. 2017;46(2). doi:10.1016/j.immuni.2017.01.007

32. Hamilton JA, Anderson GP. Mini ReviewGM-CSF Biology. Growth Factors. 2004;22(4):225–231. doi:10.1080/08977190412331279881

33. Tugues S, Amorim A, Spath S, et al. Graft-versus-host disease, but not graft-versus-leukemia immunity, is mediated by GM-CSF–licensed myeloid cells. Sci Transl Med. 2018;10(469):eaat8410. doi:10.1126/scitranslmed.aat8410

34. Lu L, Kuroishi T, Tanaka Y, Furukawa M, Nochi T, Sugawara S. Differential expression of CD11c defines two types of tissue-resident macrophages with different origins in steady-state salivary glands. Scientific Reports 2022 12:1. 2022;12(1):1–14. doi:10.1038/s41598-022-04941-5

35. McKendrick JG, Jones GR, Elder SS, et al. CSF1R-dependent macrophages in the salivary gland are essential for epithelial regeneration after radiation-induced injury. Sci Immunol. 2023;8(89):eadd4374. doi:10.1126/sciimmunol.add4374

36. de Boer J, Williams A, Skavdis G, et al. Transgenic mice with hematopoietic and lymphoid specific expression of Cre. Eur J Immunol. 2003;33(2):314–325. 10.1002/immu.200310005

37. Zubeidat K, Jaber Y, Saba Y, et al. Microbiota-dependent and -independent postnatal development of salivary immunity. Cell Rep. 2023;42(1):111981. doi:10.1016/J.CELREP.2022.111981

38. Koren N, Zubeidat K, Saba Y, et al. Maturation of the neonatal oral mucosa involves unique epithelium-microbiota interactions. Cell Host Microbe. 2021;29(2):197–209.

39. Zubeidat K, Hovav AH. Shaped by the epithelium – postnatal immune mechanisms of oral homeostasis. Trends Immunol. 2021;42(7):622–634. doi:10.1016/J.IT.2021.05.006

40. Kim M, Galan C, Hill AA, et al. Critical Role for the Microbiota in CX3CR1+ Intestinal Mononuclear Phagocyte Regulation of Intestinal T Cell Responses. Immunity. 2018;49(1):151–163.e5. 10.1016/j.immuni.2018.05.009

41. Scott NA, Mann ER. Regulation of mononuclear phagocyte function by the microbiota at mucosal sites. Immunology. 2020;159(1):26–38. 10.1111/imm.13155

42. Liu Z, Gu Y, Chakarov S, et al. Fate Mapping via Ms4a3-Expression History Traces Monocyte-Derived Cells. Cell. 2019;178(6):1509–1525.e19. 10.1016/j.cell.2019.08.009

43. Trzebanski S, Kim JS, Larossi N, et al. Classical monocyte ontogeny dictates their functions and fates as tissue macrophages. Immunity. 2024;57(6):1225–1242.e6. 10.1016/j.immuni.2024.04.019

44. Chakarov S, Lim HY, Tan L, et al. Two distinct interstitial macrophage populations coexist across tissues in specific subtissular niches. Science (1979). 2019;363(6432). doi:10.1126/SCIENCE.AAU0964

45. Dawson CA, Pal B, Vaillant F, et al. Tissue-resident ductal macrophages survey the mammary epithelium and facilitate tissue remodelling. Nat Cell Biol. 2020;22(5):546–558. doi:10.1038/s41556-020-0505-0

46. Gautier EL, Shay T, Miller J, et al. Gene-expression profiles and transcriptional regulatory pathways that underlie the identity and diversity of mouse tissue macrophages. Nat Immunol. 2012;13(11):1118–1128. doi:10.1038/ni.2419

47. Miller JC, Brown BD, Shay T, et al. Deciphering the transcriptional network of the dendritic cell lineage. Nat Immunol. 2012;13(9):888–899. doi:10.1038/ni.2370

48. Brook I. The Bacteriology of Salivary Gland Infections. Oral Maxillofac Surg Clin North Am. 2009;21(3):269–274. 10.1016/j.coms.2009.05.001

49. Liu Z, Wang H, Li Z, et al. Dendritic cell type 3 arises from Ly6C+ monocyte-dendritic cell progenitors. Immunity. 2023;56(8):1761–1777.e6. 10.1016/j.immuni.2023.07.001

50. Black S, Phillips D, Hickey JW, et al. CODEX multiplexed tissue imaging with DNA-conjugated antibodies. Nat Protoc. 2021;16(8):3802–3835. doi:10.1038/s41596-021-00556-8

51. Efremova M, Vento-Tormo M, Teichmann SA, Vento-Tormo R. CellPhoneDB: inferring cell–cell communication from combined expression of multi-subunit ligand– receptor complexes. Nat Protoc. 2020;15(4):1484–1506. doi:10.1038/s41596-020-0292-x

52. Tsou AM, Yano H, Parkhurst CN, et al. Neuropeptide regulation of non-redundant ILC2 responses at barrier surfaces. Nature. 2022;611(7937):787–793. doi:10.1038/s41586-022-05297-6

53. Jarick KJ, Topczewska PM, Jakob MO, et al. Non-redundant functions of group 2 innate lymphoid cells. Nature. 2022;611(7937):794–800. doi:10.1038/s41586-022-05395-5

54. Hauser BR, Aure MH, Kelly MC, Hoffman MP, Chibly AM. Generation of a Single-Cell RNAseq Atlas of Murine Salivary Gland Development. iScience. 2020;23(12):101838. doi:10.1016/J.ISCI.2020.101838

55. Lu CP, Polak L, Rocha AS, et al. Identification of Stem Cell Populations in Sweat Glands and Ducts Reveals Roles in Homeostasis and Wound Repair. Cell. 2012;150(1):136–150. 10.1016/j.cell.2012.04.045

56. Mauduit O, Delcroix V, Wong A, et al. A closer look into the cellular and molecular biology of myoepithelial cells across various exocrine glands. Ocul Surf. 2024;31:63–80. 10.1016/j.jtos.2023.12.003

57. Thiemann RF, Varney S, Moskwa N, Lamar J, Larsen M, LaFlamme SE. Regulation of myoepithelial differentiation. PLoS One. 2022;17(5):e0268668. 10.1371/journal.pone.0268668

58. Cansever D, Petrova E, Krishnarajah S, et al. Lactation-associated macrophages exist in murine mammary tissue and human milk. Nat Immunol. 2023;24(7):1098–1109. doi:10.1038/s41590-023-01530-0

59. Becher B, Schlitzer A, Chen J, et al. High-dimensional analysis of the murine myeloid cell system. Nat Immunol. 2014;15(12):1181–1189. doi:10.1038/ni.3006

60. Song EAC, Min S, Oyelakin A, et al. Genetic and scRNA-seq Analysis Reveals Distinct Cell Populations that Contribute to Salivary Gland Development and Maintenance. Sci Rep. 2018;8(1):14043. doi:10.1038/s41598-018-32343-z

61. Tucker AS. Salivary gland development. Semin Cell Dev Biol. 2007;18(2):237–244. 10.1016/j.semcdb.2007.01.006

62. Gao J, Li A, Fujii S, et al. p130Cas is required for androgen-dependent postnatal development regulation of submandibular glands. Sci Rep. 2023;13(1):5144. doi:10.1038/s41598-023-32390-1

63. Mellman I, Steinman RM. Dendritic Cells: Specialized and Regulated Antigen Processing Machines. Cell. 2001;106(3):255–258. doi:10.1016/S0092-8674(01)00449-4

64. Hashimoto D, Miller J, Merad M. Dendritic Cell and Macrophage Heterogeneity In&#xa0;Vivo. Immunity. 2011;35(3):323–335. doi:10.1016/j.immuni.2011.09.007

65. Haimon Z, Frumer GR, Kim JS, et al. Cognate microglia–T cell interactions shape the functional regulatory T cell pool in experimental autoimmune encephalomyelitis pathology. Nat Immunol. 2022;23(12):1749–1762. doi:10.1038/s41590-022-01360-6

66. Lazarov T, Juarez-Carreño S, Cox N, Geissmann F. Physiology and diseases of tissue-resident macrophages. Nature. 2023;618(7966):698-707. doi:10.1038/s41586-023-06002-x

67. Cabeza-Cabrerizo M, Cardoso A, Minutti CM, Pereira da Costa M, Reis e Sousa C. Dendritic Cells Revisited. Annu Rev Immunol. 2021;39(1):131–166. doi:10.1146/annurev-immunol-061020-053707

68. Guilliams M, Scott CL. Liver macrophages in health and disease. Immunity. 2022;55(9):1515–1529. 10.1016/j.immuni.2022.08.002

69. Yasuhara R, Kang S, Irié T, et al. Role of Snai2 and Notch signaling in salivary gland myoepithelial cell fate. Laboratory Investigation. 2022;102(11):1245–1256. doi:10.1038/s41374-022-00814-7

70. Fan Q, Yan R, Li Y, et al. Exploring Immune Cell Diversity in the Lacrimal Glands of Healthy Mice: A Single-Cell RNA-Sequencing Atlas. Int J Mol Sci. 2024;25(2). doi:10.3390/ijms25021208

71. Satpathy AT, KC W, Albring JC, et al. Zbtb46 expression distinguishes classical dendritic cells and their committed progenitors from other immune lineages. Journal of Experimental Medicine. 2012;209(6):1135–1152. doi:10.1084/jem.20120030

72. Yasuhara R, Kang S, Tokumasu R, Mishima K. Isolation and Functional Analysis of Myoepithelial Cells from Adult Mouse Submandibular Glands BT - Stem Cells and Lineage Commitment: Methods and Protocols. In: Turksen K, ed. Springer US; 2024:53–64. doi:10.1007/7651_2022_472

73. Bray NL, Pimentel H, Melsted P, Pachter L. Near-optimal probabilistic RNA-seq quantification. Nat Biotechnol. 2016;34(5):525–527. doi:10.1038/nbt.3519

74. Soneson C, Love MI, Robinson MD. Differential analyses for RNA-seq: transcript-level estimates improve gene-level inferences [version 2; peer review: 2 approved]. F1000Res. 2016;4(1521). doi:10.12688/f1000research.7563.2

75. Durinck S, Spellman PT, Birney E, Huber W. Mapping identifiers for the integration of genomic datasets with the R/Bioconductor package biomaRt. Nat Protoc. 2009;4(8):1184–1191. doi:10.1038/nprot.2009.97

76. Chen Y, Chen L, Lun ATL, Baldoni PL, Smyth GK. edgeR 4.0: powerful differential analysis of sequencing data with expanded functionality and improved support for small counts and larger datasets. bioRxiv. Published online January 1, 2024:2024.01.21.576131. doi:10.1101/2024.01.21.576131

77. Decoene I, Herpelinck T, Geris L, Luyten FP, Papantoniou I. Engineering bone-forming callus organoid implants in a xenogeneic-free differentiation medium. Frontiers in Chemical Engineering. 2022;4. https://www.frontiersin.org/journals/chemical-engineering/articles/10.3389/fceng.2022.892190

78. Hao Y, Stuart T, Kowalski MH, et al. Dictionary learning for integrative, multimodal and scalable single-cell analysis. Nat Biotechnol. 2024;42(2):293–304. doi:10.1038/s41587-023-01767-y

79. Korsunsky I, Millard N, Fan J, et al. Fast, sensitive and accurate integration of single-cell data with Harmony. Nat Methods. 2019;16(12):1289–1296. doi:10.1038/s41592-019-0619-0

80. Schaum N, Karkanias J, Neff NF, et al. Single-cell transcriptomics of 20 mouse organs creates a Tabula Muris. Nature. 2018;562(7727):367–372. doi:10.1038/s41586-018-0590-4

81. Kang JB, Nathan A, Weinand K, et al. Efficient and precise single-cell reference atlas mapping with Symphony. Nat Commun. 2021;12(1):5890. doi:10.1038/s41467-021-25957-x

82. Cotet TS, Agrafiotis A, Kreiner V, et al. ePlatypus: an ecosystem for computational analysis of immunogenomics data. Bioinformatics. 2023;39(9):btad553. doi:10.1093/bioinformatics/btad553

83. Andreatta M, Carmona SJ. STACAS: Sub-Type Anchor Correction for Alignment in Seurat to integrate single-cell RNA-seq data. Bioinformatics. 2021;37(6):882–884. doi:10.1093/bioinformatics/btaa755

